# Incorporating evolutionary and threat processes into crop wild relatives conservation

**DOI:** 10.1101/2021.06.15.448560

**Authors:** Wolke Tobón-Niedfeldt, Alicia Mastretta-Yanes, Tania Urquiza-Haas, Bárbara Goettsch, Angela P. Cuervo-Robayo, Esmeralda Urquiza-Haas, M. Andrea Orjuela-R., Francisca Acevedo Gasman, Oswaldo Oliveros-Galindo, Caroline Burgeff, Diana Rivera-Rodríguez, José de Jesús Sánchez González, Jesús Alarcón-Guerrero, José Sarukhán, Patricia Koleff

## Abstract

Biodiversity conservation calls for spatial explicit approaches to maximize the representation and persistence of genetic diversity given species’ idiosyncratic threats in mosaic landscapes, but conservation planning methodologies seldom account for this. Here, we introduce a novel approach that uses proxies of genetic diversity to identify conservation areas, applying systematic conservation planning tools to produce hierarchical prioritizations of the landscape. It accounts for: (i) evolutionary processes, including historical and environmental drivers of genetic diversity, and (ii) threat processes, considering taxa specific tolerance to human-modified habitats and their extinction risk status. We illustrate our approach with crop wild relatives (CWR) because their intra- and interspecific diversity is important for crop breeding and food security. Although we focus on Mesoamerican CWR within Mexico, our methodology offers new opportunities to effectively guide conservation and monitoring strategies to safeguard the evolutionary resilience of any taxa, including in regions of complex evolutionary histories and mosaic landscapes.

## Introduction

The Anthropocene has driven biodiversity to persist in mosaic landscapes, imposing new challenges for conservation. Actions to protect diversity at the genetic level have been limited, though crucial for evolutionary resilience.^1^ Systematic conservation planning (SCP) can aid the process of effectively locating and managing areas to protect biodiversity. However, most SCP assessments focus on biodiversity patterns rather than persistence (specie’s capacity to continue existing in an area), and do not incorporate ecological and evolutionary processes nor the distribution of genetic diversity.^1–4^ This is particularly urgent for crop wild relatives (CWR; wild plants closely related to crops) which are at the crossroads between biodiversity conservation and food security, given the importance of their genetic diversity to adapt crops to challenging environments.^5^

As in any wild species, CWR genetic diversity depends on demographic history, population structure and natural selection,^11,12^ which changes according to the climatic and geologic history of a given area.^13^ Most regions where crops were originally domesticated (centres of domestication)-thus often holding high diversity of CWR- are tropical or topographically heterogeneous,^14^ promoting long-term population persistence and genetic differentiation in complex patterns.^15,16^ To address the processes behind CWR genetic diversity, SCP should consider both current and historical drivers of evolution. Another challenge for SCP is to account for CWR interaction with agriculture. Most CWR are threatened by land use change, but some thrive in agricultural landscapes and are tolerated or fostered by farmers.^6–10^ Additionally, in some regions gene flow between crops and their CWR can occur, and has been part of the crop domestication process for thousands of years.^9,10,17,18^ However, in modern agriculture, which is linked to the cultivation of highly genetically uniform commercial varieties and living modified organisms, gene flow can become a risk in the form of genetic assimilation (crop alleles replacing wild ones) and demographic swamping (wild populations shrink due to hybridization).^19^

Substantial advances in CWR conservation planning have been done based on Magos Brehm et al.^20,21^ methodology. Several countries have adopted this workflow (e.g.^22–24^), greatly contributing to CWR conservation globally^25^. Notwithstanding, most CWR planning apply the minimum set cover problem to detect the least number of sites to complement existing reserves, while representing genetic diversity with ecogeographic variables. Although useful in some cases,^20,25^ this approach is not fully suitable to capture, in a representative way, the genetic diversity spectrum present in centers of origin and domestication, such as Mexico, where genetic diversity is driven by complex climatic, geologic and human histories,^26^ and where crops and their CWR co-exist.^27–29^

Here, we introduce a novel framework to identify conservation areas based on a hierarchical prioritization of the landscape, to account for: (i) evolutionary processes, including historical and environmental drivers of genetic diversity (hereafter, proxies of genetic diversity, PGD) and (ii) threat processes, considering taxon specific tolerance to human-modified habitats and IUCN extinction risk status.

## Results

In the context of the project ‘Safeguarding Mesoamerican crop wild relatives’, a collaborative partnership among government agencies, local communities, universities and NGOs from El Salvador, Guatemala, Honduras, Mexico, the UK and IUCN, we followed the main steps of Magos Brehm et al.^20,21^ workflow: (i) CWR checklist, (ii) CWR inventory, i.e. subsetting the CWR checklist, (iii) taxa extinction risk assessment, and (iv) spatial analysis for *in situ* and *ex situ* conservation, introducing a novel approach to account for genetic variation (Fig. 1). For the SCP analyses, we used the Zonation software,^30,31^ incorporating potential species distribution models (SDM) divided by PGD, and taxa habitat preference maps to identify suitable locations with reference to the vegetation and land use and land cover map.

**Fig. 1.**
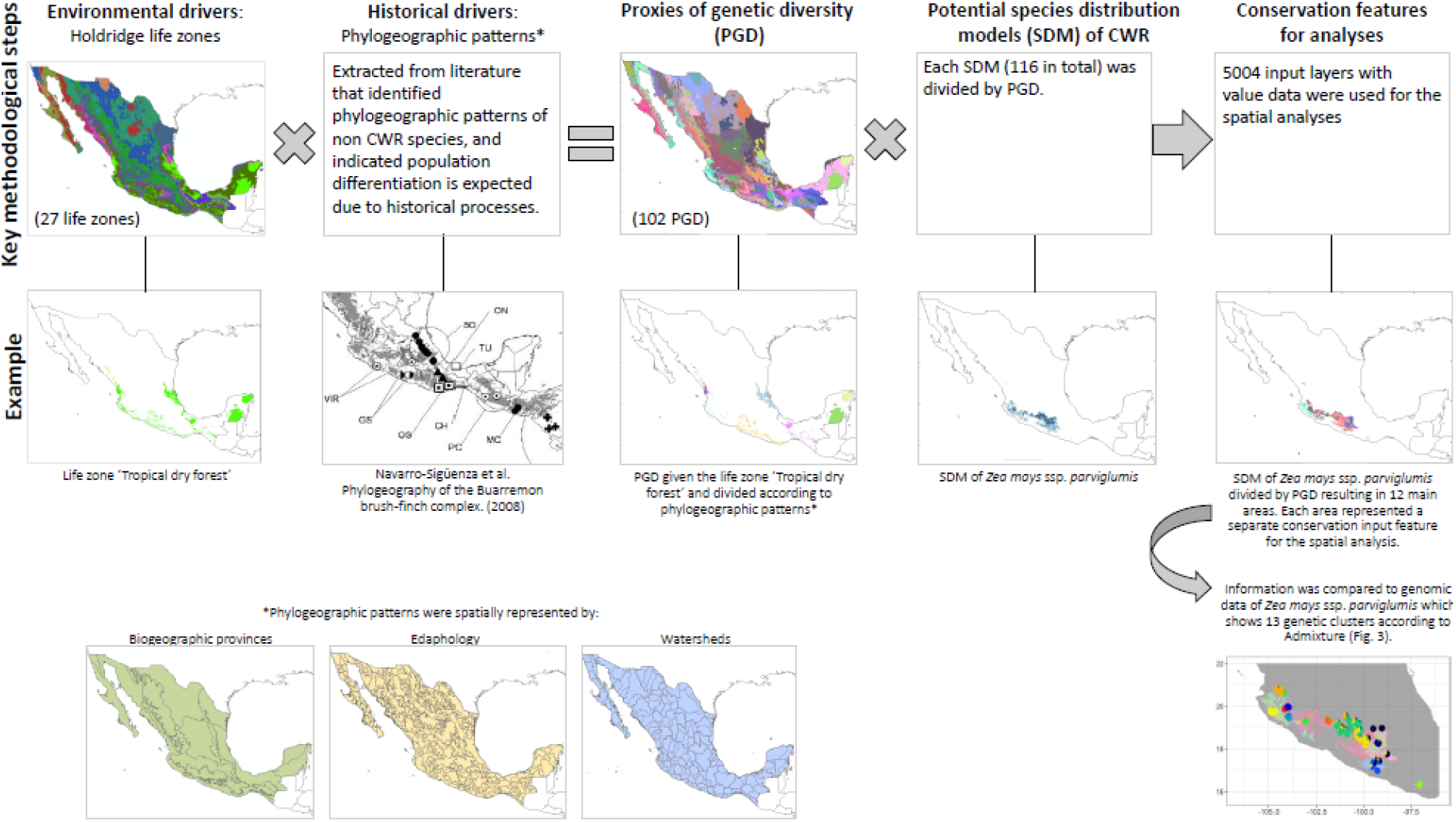
Methodological framework to develop proxies of genetic diversity to assess conservation areas in the context of the project “Safeguarding Mesoamerican crop wild relatives”. (See Supplementary Information for details.)

The first three steps are detailed in Goettsch *et al*.^8^ Here, we used the inventory of 224 native or endemic species, subspecies, varieties and races of Mesoamerican CWR, related to nine crops: chili pepper (*Capsicum* spp.), squash (*Cucurbita* spp.), cotton, (*Gossypium* spp.), avocado (*Persea* spp.), bean (*Phaseolus* spp.), husk tomato (*Physalis* spp.), potato (*Solanum* sect. *Petota*), maize *(Zea* spp., and *Tripsacum* spp.), and vanilla (*Vanilla* spp.) (Table 1, Supplementary Table 3). Of these, 74 of the 310 taxa were previously identified as global priority CWR for Mexico.^22^ These taxa were comprehensively assessed for the IUCN Red List of Threatened Species, resulting in seven critically endangered, 47 endangered, 16 vulnerable and nine near threatened taxa (Goettsch *et al*.^8^).

**Table 1.** Mesoamerican crop wild relatives selected for extinction risk assessment.

| Crop common name | CWR genus | No. of taxa | IUCN Red List Category |  |  |  |  |  | No. of taxa with potential species distribution model |
| --- | --- | --- | --- | --- | --- | --- | --- | --- | --- |
|  |  |  | Critically endangered (CR) | Endangered (EN) | Vulnerable (VU) | Near threatened (NT) | Least concern (LC) | Data deficient (DD) |  |
| Chili pepper | *Capsicum* | 4 |  | 1 |  |  | 3 |  | 4 |
| Squash | *Cucurbita* | 11 |  | 1 |  | 2 | 5 | 3 | 10 |
| Cotton | *Gossypium* | 13 | 2 | 6 | 4 |  | 1 |  | 6 |
| Avocado | *Persea* | 18 |  | 7 | 2 | 1 | 5 | 3 | 12 |
| Bean | *Phaseolus* | 55 |  | 12 | 5 |  | 36 | 2 | 22 |
| Husk tomato | *Physalis* | 67 | 2 | 3 | 2 | 4 | 47 | 9 | 30 |
| Potato | *Solanum* | 26 |  | 5 | 1 | 2 | 16 | 2 | 18 |
| Maize | *Tripsacum* | 11 |  | 3 | 1 |  | 7 |  | 3 |
| Maize | *Zea* | 11 | 2 | 3 | 1 |  | 5 |  | 9 |
| Vanilla | *Vanilla* | 8 | 1 | 6 |  |  |  | 1 | 2 |
| **Total** | **10** | **224** | **7** | **47** | **16** | **9** | **125** | **20** | **116** |

Based on potential SDM of Mesoamerican CWR, areas of high taxa richness were identified along the Trans-Mexican Volcanic Belt, and in the montane areas of Oaxaca and Chiapas in Southern Mexico (Fig. 2). The spatial distribution pattern was consistent with global richness of CWR^32^ and trends in other taxonomic groups showing higher species richness in heterogeneous and montane environments.^33,34^ The global study highlighted montane areas of Mexico as showing high CWR richness with around 35 taxa.^32^ Our results show more taxa per area than the previous study (>50 taxa of CWR by square km; Fig. 2; Supplementary Fig. 1), despite using only 10% of taxa and using potential distribution models, which were validated by experts and independent data. Other CWR analysis for Mexico estimated up to 167 taxa in a 15×15 km grid square.^22^

**Fig. 2.**
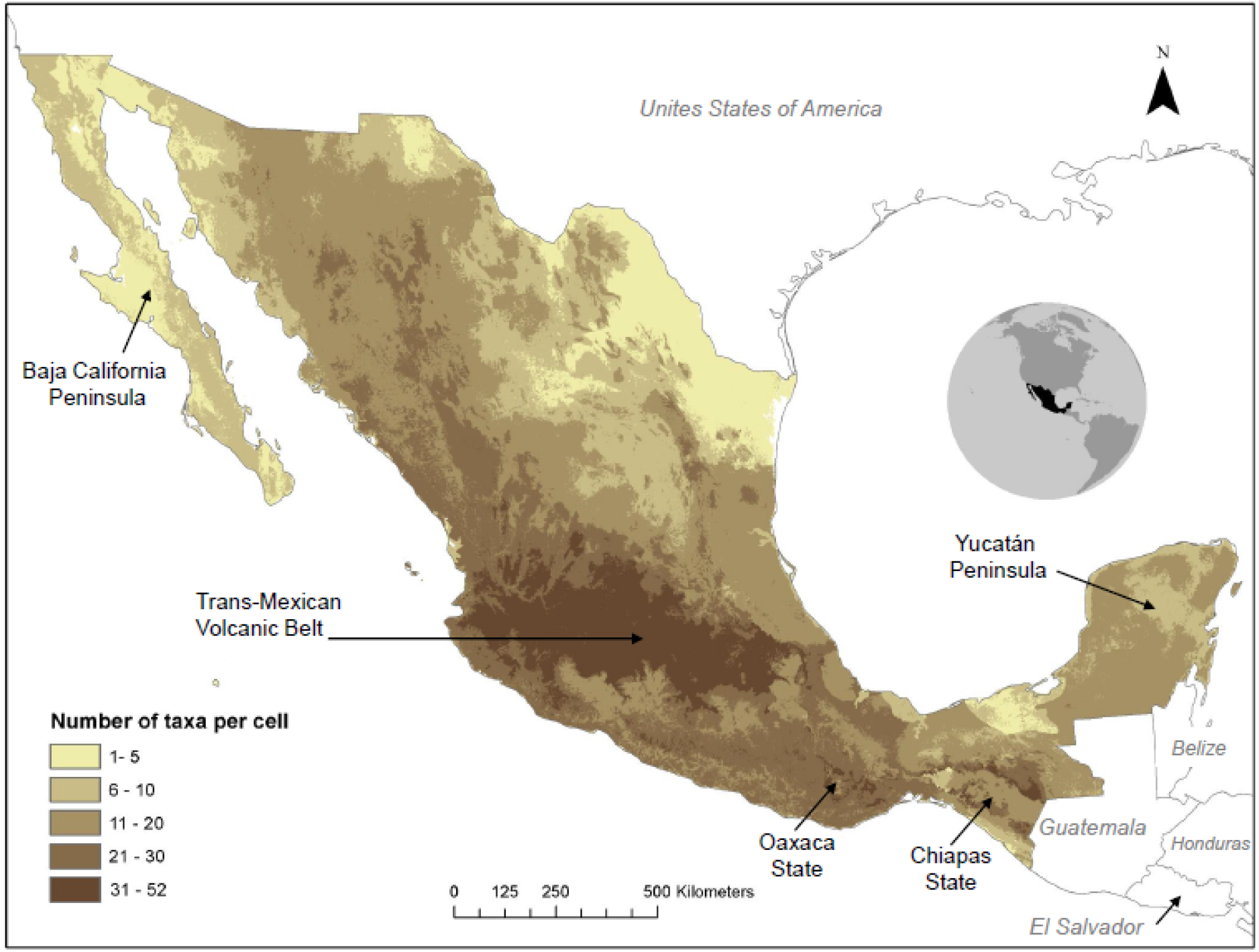
Estimated crop wild relatives richness in Mexico, based on 116 taxa with potential species distribution models (see Supplementary Information for details). Spatial resolution 1 km^2^.

To identify conservation areas, we used the Zonation software,^30,31^ to establish a hierarchical prioritization of the landscape and to optimize the representation of taxa or other conservation features in a given area, by incorporating taxa specific information (Supplementary Information 1). Critical to this approach was the development of PGD to reveal genetic differentiation in the absence of genomic information. We delimited PGD based on Holdridges’ life zone classification, characterized by specific ranges of temperature, humidity and potential evapotranspiration (Supplementary Table 5, Supplementary Fig. 2). Then each life zone was divided according to phylogeographic patterns of non CWR species, using biogeographic regions or topographic and edaphic data to define the cutline (see Fig. 1; Supplementary Table 6; Supplementary Fig. 3). The first step is similar to the ecogeographic land characterization proposed by Parra-Quijano et al.,^35^ that assumes that adaptive genetic features vary according to environmental variation. Additionally, the present approach incorporates areas where population differentiation is expected due to historical processes, which is particularly important in tropical and topographically complex areas.^26^

**Fig. 3.**
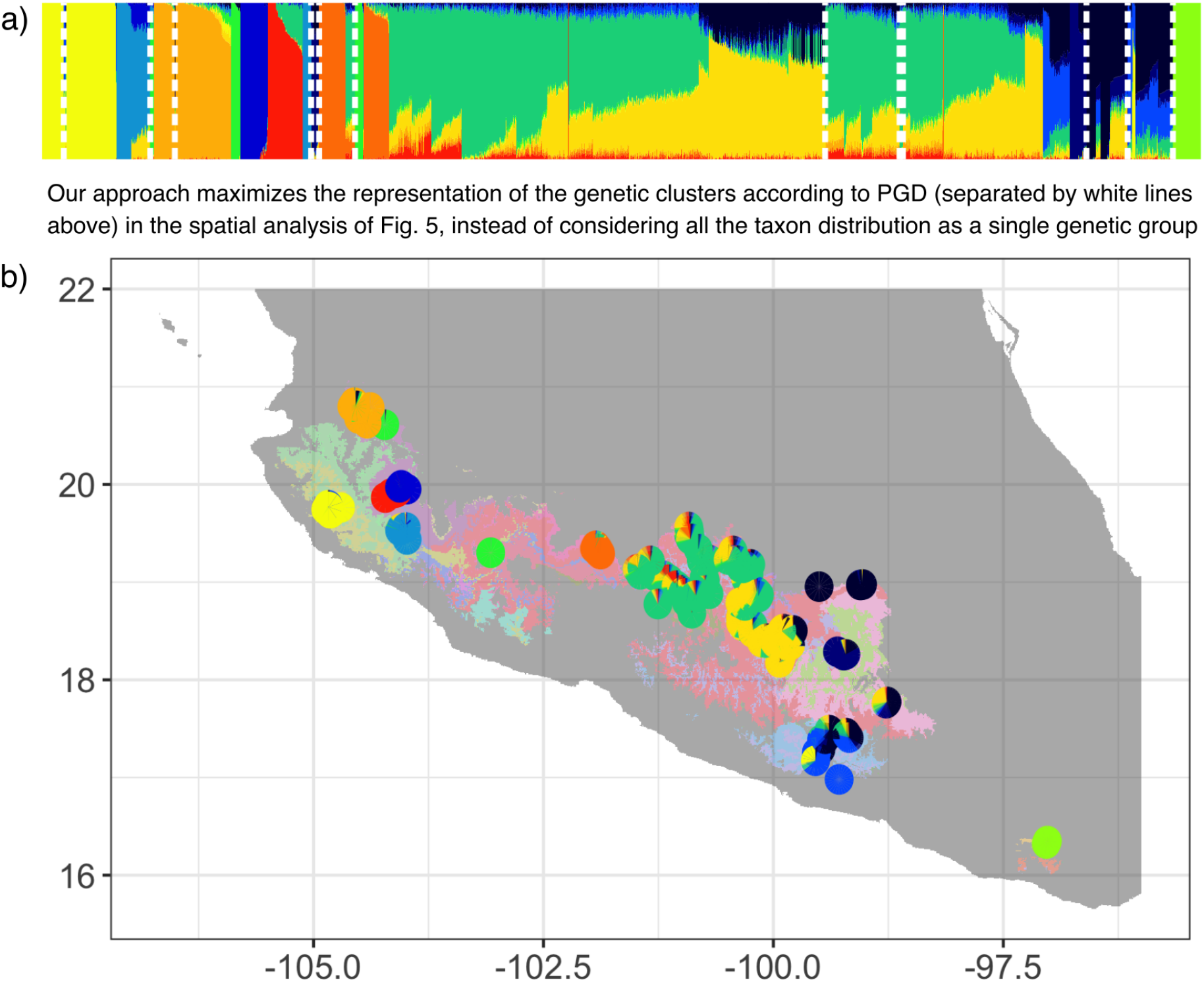
Genetic diversity of *Zea mays* ssp. *parviglumis* represented in the proxies of genetic diversity (PGD). a) Admixture plot assuming K=13 genetic clusters, using ca. 30,000 SNPs (data from Rivera-Rodríguez^48^). Each bar represents the proportion of different genetic clusters (colours) conforming an individual. White dashed lines represent the PGD where the samples fell. b) Potential species distribution model of *Z. mays* ssp. *parviglumis* as given by PGD (background colours) and proportion of genetic clusters by sampling locality (pie plots). Colours of genetic clusters in (a) and (b) match. Spatial resolution 1 km^2^.

Our PGD were supported by available empirical data from the teosinte *Zea mays* ssp. *parviglumis*. According to a population analysis with >30,000 single nucleotide polymorphisms (SNPs) data, *Z. mays* ssp. *parviglumis* is structured in 13 genetic groups. Normally, SCP would use SDM without differentiating these genetic clusters within the taxon distribution range, but our approach allows maximizing their representation in the spatial solution (Fig. 3; Supplementary Fig. 4). While there is no complete coincidence between the observed and estimated number of populations, using PGD allowed to incorporate, at least partially, potential population genetic differentiation in a spatially explicit manner. PGD can be assessed for any taxa without genetic data, which is particularly important if we aim to secure genetic variation within all taxa.^36^ As genomic resources become available for more species, the PGD could be further corroborated and fine-tuned.

**Fig 4.**
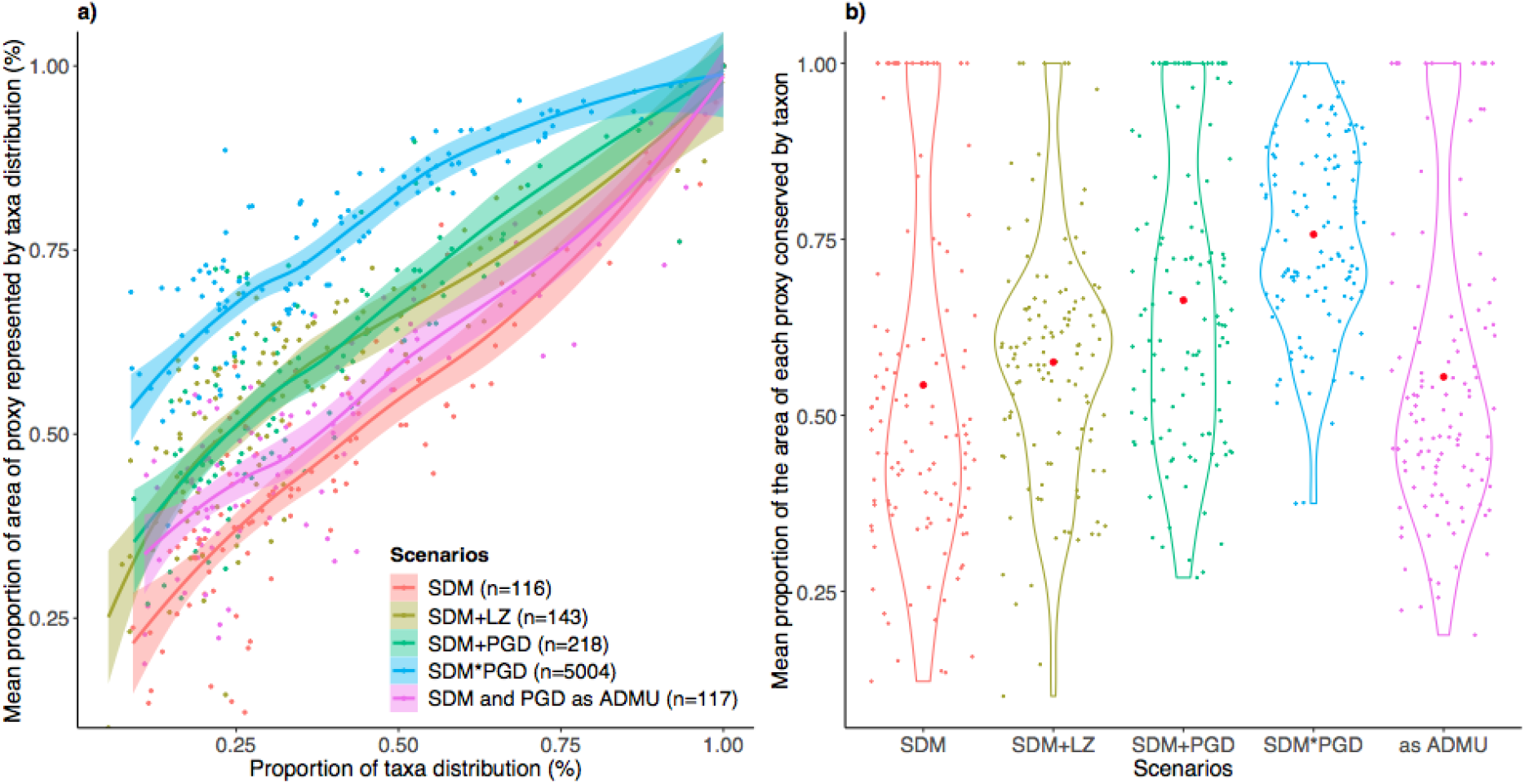
Performance of five scenarios to represent conservation features of Mesoamerican crop wild relatives, considering 20% of Mexico’s terrestrial area. a) Representation curves of the proportion of taxa distributions and mean proportion of proxies of genetic diversity (PGD) areas within them; b) violin plots showing the mean proportion of the area of each PGD represented within each taxon. Scenarios: SDM - considered 116 taxa with potential species distribution models; SDM+LZ - considered 116 taxa with potential species distribution models and 27 life zones; SDM+PGD - considered 116 taxa with potential species distribution models and 102 PGD; SDM*PGD - considered 5004 input layers, that resulted by subdividing each potential species distribution model by PGD; SDM and PGD as ADMU - considered 116 taxa with potential species distribution models as conservation features and 102 PGD as administrative units.

To show the importance of explicitly accounting for genetic variation in SCP, we tested different alternatives to incorporate PGD into the spatial analysis, and compared it against results excluding them, or using only Holdridge’s life zones (Fig. 4). The approach that maximized the representation of both taxa and PGD, within the taxa potential distribution ranges (hereafter taxa ranges), subdivided each potential distribution model by PGD, resulting in 5,004 input layers for the spatial analysis (SDM*PGD scenario). With this, on average 41% of each taxon range and 76% of the area of each PGD per taxon were represented when evaluating 20% of the country. Other scenarios also captured the potential genetic variation inferred through the use of proxies, but were less efficient. When considering the potential distribution models and PGD (SDM+PGD scenario), or using PGD as the unit of analysis (SDM and PGD as ADMU scenario), on average 37% and 48% of the range of each taxon was represented in 20% of the country, respectively, but the average area of each PGD within each taxon range was smaller compared to other scenarios (SDM scenario: 57%; SDM and PGD as ADMU scenario: 66%; SDM*PGD scenario: 76%). Performing the analysis only with potential distribution models (SDM scenario), resulted in the highest proportion of area of taxa ranges (on average 48%), but the poorest representation of PGD within the area of each taxa (on average 54%). This suggests that performing SCP analyses without explicitly considering an indicator for genetic variation fails to represent the heterogeneity within the taxa ranges, and thus likely its associated genetic diversity. Since both UN Sustainable Development Goal 2.5 ^37^ and Aichi Target 13 of CBD call to maintain genetic diversity within domesticated plants and wild relatives, it is particularly urgent for SCP to specifically incorporate it. By incorporating indicators of genetic variation, our approach contributes to focus on conservation measures across areas that together enhance species evolutionary resilience in the face of climate change and other threats, which was recognised as a priority more than a decade ago.^1^ Additionally, PGD-restricted taxa, i.e. revealing a taxon specific attribute related to its genetic diversity representation, should also receive high attention for both *in situ* and *ex situ* conservation actions.

For the final SCP analysis, we incorporated: (a) the CWR potential distribution models subdivided by PGD, so potential populations of specific and infraspecific levels could be recognized as different features in the spatial assessment; (b) the IUCN threat categories to weight taxa (giving higher values to taxa at higher risk of extinction; Supplementary Table 3); (c) occurrence records for taxa without potential distribution model, to ensure their representation in the solution (Supplementary Table 3); and (d) taxon specific habitat preferences to identify suitable locations considering land-use and land-cover, including different types of agricultural systems (Supplementary Table 7; Supplementary Figure 5).

**Fig. 5.**
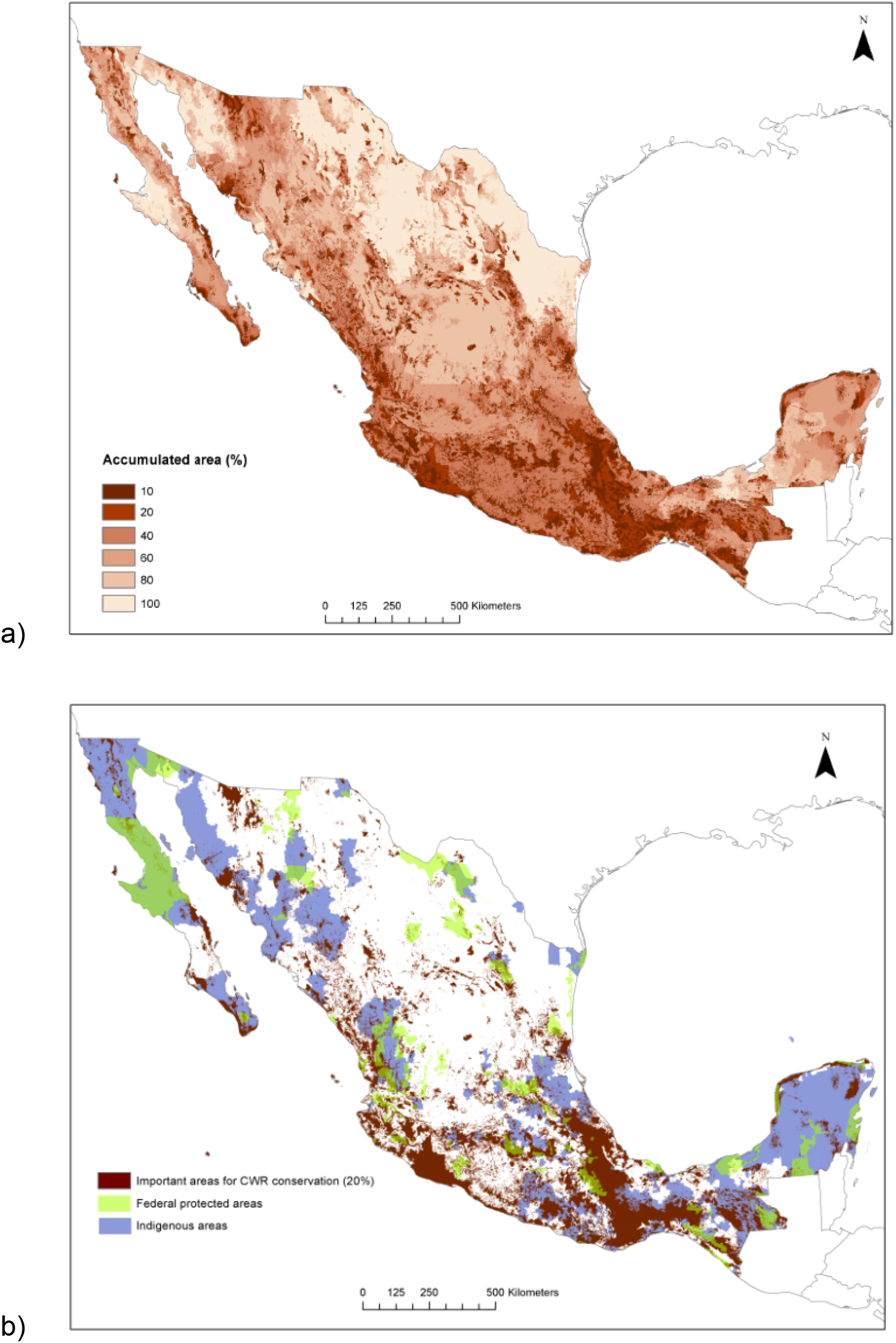
Results of the systematic conservation planning process for Mesoamerican crop wild relatives in Mexico, considering potential species distribution models subdivided by proxies of genetic diversity (PGD; see text for details). a) Continuous map; b) Conservation area proposal considering 20% of Mexico’s terrestrial area, highlighting federal protected areas and indigenous areas. Spatial resolution 1 km^2^.

CBD Aichi Target 11 encouraged parties to protect at least 17% of terrestrial regions^32^, therefore, we examined the solution considering 20% of Mexico (Fig. 5). The identified conservation areas are located in the temperate mountain areas of the Trans-Mexican Volcanic Belt, characterized by high species richness and endemism; in the region from central Veracruz state to Chiapas, crossing Puebla and including large areas of Oaxaca, the Tehuacán-Cuicatlán valley and the Chimalapas region, corresponding to areas of highly heterogeneous environments; along the northern coastline of the Michoacán state in Central Mexico, and in the cloud forests and rainforests of southern Mexico, representing habitats for range-restricted species such as *Vanilla odorata*; and in the arid and semi-arid states of Sonora and Baja California, were *Gossypium* species are common (Fig. 5 a,b). Almost half of these important areas for conservation were located within indigenous areas and 11% were located within federal protected areas (PAs) (Fig. 5b). Also, one third of Areas Voluntarily Destined for Conservation (ADVC, by its Spanish acronym) are covered in the selected 20% area. PAs represent great opportunities for active conservation and implementation of management and monitoring of CWR^39^, but because ADVC are a biocultural approach for sustainable management; they in particular offer an opportunity to further support CWR conservation. For all kinds of PAs, a needed step to promote CWR conservation in management plans is to generate comprehensive inventories of CWR occurring within them, ^8,40^ although other conservation approaches should be considered too.^8^

Performance curves showed the effectiveness of the spatial solution at representing CWR potential distribution ranges and its genetic diversity. For instance, in 20% of the country, on average, 50% of the area of each PGD of each taxa were represented (Fig. 6; Supplementary Fig. 9). Representation values of taxa grouped by threat categories differed due to the conservation weights established according to their extinction risk. Noteworthy, it was impossible to represent 100% of the taxa ranges, and consequently its PGD, revealing that for many taxa a considerable amount of their habitat has already been lost, degraded and fragmented within their potential range. This is particularly critical for the most threatened taxa as more than 40% of their potential distribution range may no longer have suitable conditions. Also, it indicates the urgent need for both conservation and restoration actions to maintain and recover natural vegetation, but might also imply that some genetically distinct populations could have already been lost. Local extinction of populations has been suggested as an important indicator of loss of genetic diversity.^36^

**Fig. 6.**
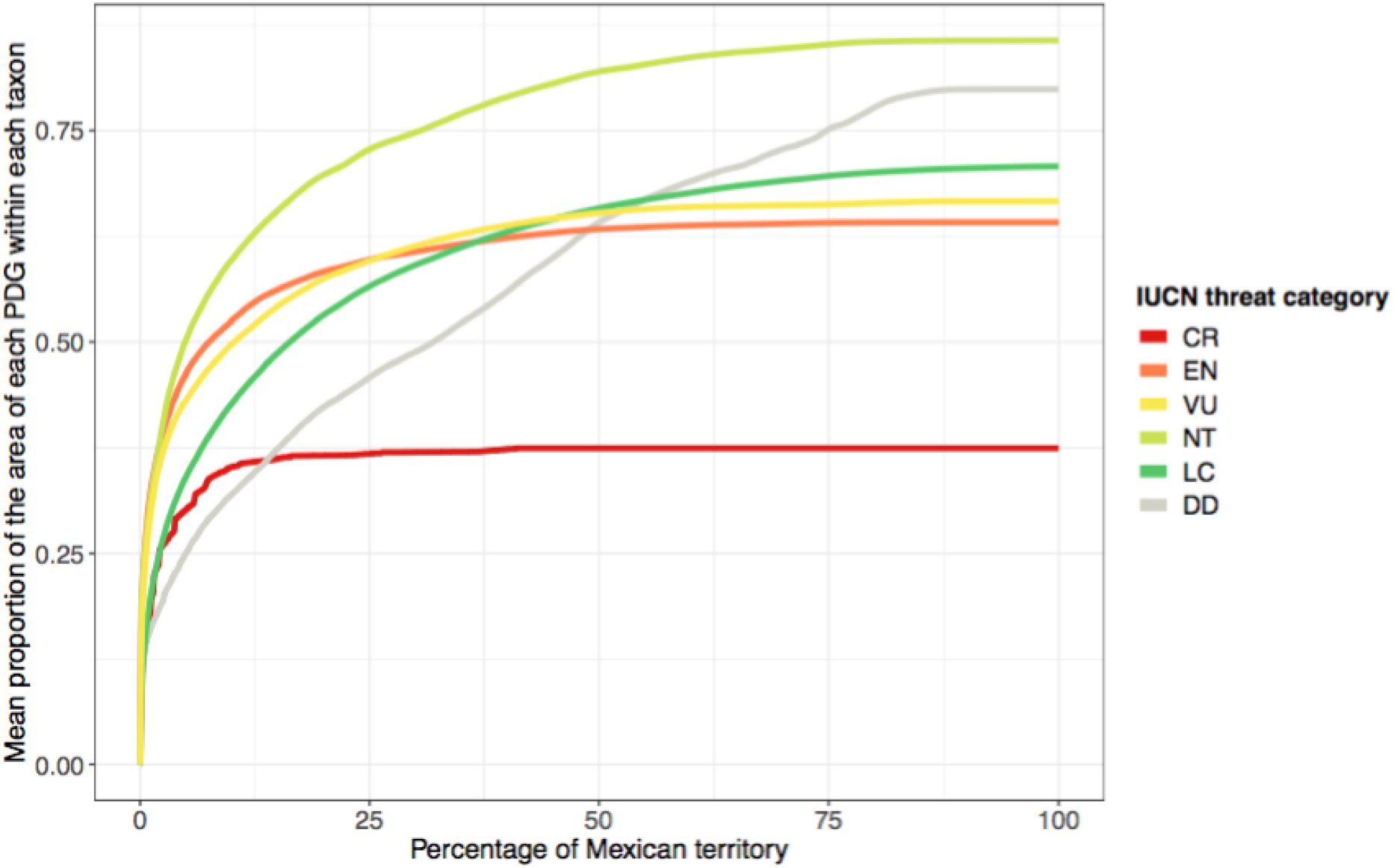
Performance curves quantifying the proportion of crop wild relatives within the scenario considering potential species distribution models subdivided by proxies of genetic diversity (PGD; see text for details). Data is grouped by IUCN Red List Category of taxa (CR=Critically Endangered, EN=Endangered, VU=Vulnerable, NT=Near threatened, LC=Least concern, DD=Data deficient). Performance was evaluated for 116 taxa with potential species distribution model.

Although taxa might be represented at least once in a minimum number of pixels (Fig. 6), the conservation of ecological and evolutionary processes shaping biodiversity at all levels (genes, populations, species, ecosystems) can not be secured in only a fraction of cells, in particular in Mexico where plant domestication continues to occur today in the hands of millions of smallholder farmers through sustainable resource management. Therefore, the hierarchical priority rank map (Fig. 5a) and selected conservation areas (Fig. 5b) offer a guide for implementing conservation and sustainable landscape management measures at large spatial areas, e.g. landscapes, ecoregions, or basins to support biodiversity, including agroecosystems, and the services and benefits they provide.^41^ In addition, the continuous map, i.e. without setting a specific target for a certain percentage of the landscape (Fig. 5a), provides a broad overview to highlight that regional and local conservation is needed all over Mexico. Directed conservation action and sustainable resource management at landscape level across the country, acknowledging the importance of CWR populations and their interactions with domesticated species, would support critical ecological and evolutionary processes, but see Contreras-Toledo et al.^22^ for a representation approach at pixel level.

Since taxa have considerably distinct habitat preferences and life histories (Supplementary Table 7), they can be affected differently by land use change and agricultural practices^8^: eg. *Persea* are trees distributed in well-preserved vegetation, such as cloud forests, while *Gossypium* are bushes requiring a certain degree of disturbance,^43^ and *Capsicum* and *Physalis* are managed in the wild or tolerated within traditional agricultural systems.^6,7^ To identify conservation priorities for different land use preferences, we ran three scenarios, including: all taxa (Fig. 5), taxa exclusively distributed in well-preserved vegetation, and taxa that can be associated to different habitats and land uses (e.g. natural vegetation, agriculture and urban areas). Results showed different spatial patterns (Supplementary Figs. 6, 7) and performance curves (Supplementary Fig. 8), exposing the habitat preferences of taxa included in each scenario (Supplementary Fig. 10). Explicitly accounting for the effect of land use on conservation of CWR as done here can allow to promote synergic planning and actions among different sectors, specially between the environmental and the agricultural.^8^ Further analysis could also consider future land use and climate change scenarios to assess conservation priorities in the long term.

While our analysis has focused on *in situ* conservation, it could be also useful to address the challenges of *ex situ* conservation. Namely, the spatial results may indicate sampling areas based on either taxon rich or important areas of CWR conservation, i.e. where range restricted taxa distribute, in order to reduce the collection efforts, or poorly explored areas to cover representation gaps.

In summary, our approach identifies conservation areas of high CWR taxa richness and uniqueness, and represents genetic diversity in a spatially explicit way, accounting for historical and environmental drivers of genetic differentiation. The major limitation of our study is the lack of high resolution genetic data to corroborate if the PGD are reliable across different taxa and to fine-tune the approach. However, given the rate at which biodiversity is declining, it is better to include an inaccurate representation of genetic diversity within taxa, than to perform conservation assessments without explicitly accounting for it. Our approach might be challenging in terms of preparing and handling large datasets; running analysis with thousands of input layers needed to be done in a computing cluster. Still, a major benefit of the software Zonation is its ability to incorporate different sources of data (i.e. potential SDM, occurrence records, PGD, land use and cover maps, threat categories), and to link each biodiversity feature to a certain condition group, eg. habitat preference. One of the results is a hierarchical representation of the landscape that can inform proactive and reactive conservation measures. Also, it is not necessary to define area specific representation targets; sites are ranked according to occurrence levels, conservation weight of biodiversity features, and other considerations. A general advantage of our SCP approach is that it allows incorporating key information to enhance biodiversity conservation by addressing genetic diversity as well as environmental and evolutionary processes at the landscape level. Our framework can be used for the establishment of conservation areas, but also to promote sustainable management across the landscape. Thus, conservation and development goals can be tackled simultaneously in order to achieve long-term sustainability.

## Discussion

Identification of areas of high conservation value that consider the representation of genetic diversity is needed for strategic planning and decision-making; and in the case of Mesoamerica it supports the development of National Strategic Action Plans for the conservation and use of CWR. Our approach allows maximising the representation of genetic diversity and threatened and range limited taxa, while accounting for habitat preferences related to land use. Additionally, our hierarchical representation of the landscape offers a broader perspective not only to identify where area-based conservation measures are mostly required, but also to implement sustainable development policies that strengthen rural communities and economies^42^. In the case of CWR, this can inform public policy regarding living modified organisms such as crops^27^ and agriculture subsidies^44^ in order to mitigate threat processes to CWR.^8^ Incorporating SCP outputs to the design of cross-sectoral policies can allow to move *in situ* conservation of CWR beyond their protection only in PAs. This is of particular relevance for regions that are centres of origin, domestication and diversification of crops, and where sustainably managed landscapes can not only contribute to halting biodiversity loss but also to the provision of evosystem services. ^41,45,46^

We focused on CWR due to their importance for food security and adaptation of crops to changing environments.^5^ By explicitly including genetic diversity of Mexican CWR in SCP, our results and associated policy suggestions^47^ contribute to achieving Mexican commitments to UN Sustainable Development Goal 2.5 and CBD Aichi Target 13.^45^ However, as recent suggestions to the CBD post-2020 strategy highlight, we also need to conserve and monitor genetic diversity beyond domesticated species and their relatives.^36^ Our methodology could be applied to any taxa, and thus represents an important contribution to achieve this new goal at the global level.

## Methods

A detailed description of the methods and results is in the Supplementary Information 1. Supplementary tables and figures are available separately at Supplementary Information 2. Data and scripts are available at Dryad XXX and Github https://github.com/AliciaMstt/analisisUniCons_proxiGen repositories. Potential Species distribution models can be downloaded at XXX (available upon acceptance) or see Supplementary Table 9 for taxon specific links.

## Acknowledgments

We are grateful to the Darwin Initiative of the United Kingdom for providing funding, and to the International Union for Conservation of Nature, IUCN, for implementing the project “Safeguarding Mesoamerican Crop Wild Relatives” (project number: 23-007). We thank all participants who collaborated during the Darwin Initiative project: Araceli Aguilar-Meléndez, Gabriel Alejandre-Iturbide, Flavio Aragón Cuevas, César Azurdia Pérez, Jamie A. Carr, Gabriela Castellanos-Morales, José G. Cerén López, Aremi Contreras Toledo, María Eugenia Correa-Cano, Lino de la Cruz Larios, Daniel G. Debouck, Alfonso Delgado-Salinas, Manuel González-Ledesma, Enrique González-Pérez, Mariana Hernández-Apolinar, Braulio E. Herrera-Cabrera, Megan Jefferson, Rafael Lira-Saade, Francisco Lorea-Hernández, Mahinda Martínez, Jenny Menjívar, María de los Ángeles Mérida, Aura Morales, Caroline M. Pollock, Martín Quintana-Camargo, Aarón Rodríguez, José A. Ruíz Corral, Guillermo Sánchez-de la Vega, Mariella Superina, Marcel F. Tognelli, Ofelia Vargas-Ponce, Melania Vega, Ana Wegier, Pilar Zamora Tavares. Special thanks to: Richard K.B. Jenkins as the IUCN project leader; Emma Gómez Ruiz as former responsible of the project; our colleagues from CONABIO, particularly Oscar Godínez Gómez, Diana Ramírez Mejía, Juan Barrios Vargas, Ernesto Campos Murillo, Daniel Ortiz Santamaría, and Cuauhtémoc Enríquez García for data processing and technical help; Gloria Espinosa Sánchez, Patricia Galindo Hernández and Nancy Corona Pedroza for administrative support; and the great work of the General Direction of Science Outreach at CONABIO; Rosalba Becerra for her creative designs; Andrés Lira Noriega, Daniel Piñero and Jorge Soberón for feedback on the manuscript; and Shelagh Kell and Nigel Maxted for comments to our approach and their effort to conserve CWR. Analyses were carried out on CONABIO’s computing cluster, supported by Ernesto Campos Murillo and the Subcoordinación de Soporte Informático. Also, we would like to express our genuine gratitude to the experts that contribute to the knowledge of biodiversity of Mesoamerica, and to the small-scale farmers and traditional households that maintain the processes of evolution and domestication of crop landraces.

