## Supplementary Materials 1 for "Incorporating evolutionary and threat processes into crop wild relatives conservation"

### Supplementary Information 1

#### Methods

We followed the planning approach suggested by Magos Brehm et al.[1] for crop wild relatives (CWR) conservation. We addressed the main steps of the toolkit: (i) CWR checklist, (ii) CWR inventory, i.e. subsetting the CWR checklist, (iii) taxa extinction risk assessment, and (iv) spatial analysis for *in situ* and *ex situ* conservation, introducing a novel approach to account for genetic diversity (Fig. 1; see Spanish version[2]).

During the process -framed under the project "Safeguarding Mesoamerican crop wild relatives" (https://www.darwininitiative.org.uk/project/23007/)- more than 100 experts from academic, governmental, and non-governmental organizations from El Salvador, Guatemala, Honduras, Mexico, the UK, and IUCN participated in six workshops, shared data, and provided fundamental knowledge and feedback at each project stage to ensure accurate, reliable and robust information for next steps (incl. checklist, inventory, extinction risk assessment, and inputs and outputs for spatial analysis).

The checklist, inventory and risk assessment were collaboratively developed, and incorporated CWR distributed in El Salvador, Guatemala, Honduras, and Mexico, hereafter called Mesoamerica. The methodology of these first three steps is detailed in Goettsch et al.[3] Here, we describe the spatial analysis to identify areas for *in situ* and *ex situ* conservation of CWR in Mexico, which was done independently by each country.

To identify conservation areas of CWR in Mexico, we assessed proxies of genetic diversity (PGD) that account for (i) evolutionary processes, including historical and environmental drivers of genetic diversity within each taxon, and (ii) threat processes, by considering taxon specific tolerance to human-modified habitats and IUCN extinction risk category. We applied a systematic conservation planning (SCP) approach and performed spatial analysis using the software Zonation.[4] We compared different scenarios to represent genetic diversity of CWR based on potential species distribution models (SDM) and proxies of genetic diversity (PGD).

##### Study area

Mesoamerica is a cultural region encompassing the territories of Belize, Guatemala, El Salvador, the southern part of Mexico and the western region of Honduras, Nicaragua and Costa Rica. In this study, we also included the dry areas of northern Mexico that are part of Aridmerica[5] and the Nearctic biogeographic realm[6] to account for the full extent of the geographic range of many taxa included in the extinction risk assessment.[3]

For the present analysis, we focused on Mexico, which is one of the most biodiverse countries in the world.[7] Its high biological diversity is attributed to its geographic, topographic, climatic, geological and cultural characteristics, which, among other factors, shaped the distribution of an extraordinary variety of ecosystems and number of species with high levels of endemism and species turnover among different regions.[8–11] In particular, the high genetic variation within populations of landraces and CWR is the result of past and ongoing sociocultural processes occurring in a wide range of distinct environmental conditions.[12,13]

#### CWR checklist and inventory

The compiled CWR checklist included more than 3,000 species and subspecies of 92 genus and 45 families that belong to the same genus of a crop cultivated or (semi-) domesticated in Mesoamerica (Supplementary Table 1).

To subset the list, a first set of criteria were established during the first stakeholder workshop, these included: 1) occurrence of wild relatives of cultivated or domesticated taxa in Mesoamerica; 2) the existence of research groups working on the taxa that could support the extinction risk assessment; 3) related to a crop of economic and nutritional importance at local, national and regional levels, or cultivars known to require genetic improvement. At the species level, similar criteria were used, and additionally included: 4) natural distribution of CWR of a selected genus in Mesoamerica, incl. Aridamerica; 5) inclusion in the primary or secondary gene pool, and in some cases the tertiary gene pool, that is more distantly related to the taxa of the primary gene pool but which can have important adaptive traits. This preliminary grouping included CWR taxa related to avocado, cotton, amaranth, cocoa, squash, sweet potato, chayote, chili pepper, cempasuchil, bean, sunflower, maize, papaya, potato, vanilla, and yuca.

Based on these criteria, each country compiled a preliminary list of CWR of these crops, which were integrated into a single database, consisting of more than 500 taxa related to the selected crops (Supplementary Table 2). The list had to be further reduced, due to time and funding restrictions, for which the feasibility to comprehensively conduct the risk assessment of all species and subspecies of a taxonomic group was taken into account. Thus, not all species in the group necessarily met the five criteria previously mentioned.

The final Mesoamerican CWR inventory included 224 taxa of nine crops, considering chili pepper (four *Capsicum* spp.), squash (11 *Cucurbita* spp.), cotton (13 *Gossypium* spp.), avocado (18 *Persea* spp.), bean (55 *Phaseolus* spp.), husk tomato (67 *Physalis* spp.), potato (26 *Solanum* sect. *Petota* spp.), 22 taxa of maize (11 *Zea* spp. and 11 T*ripsacum* spp.), and vanilla (eight *Vanilla* spp.) (Table 1, Supplementary Table 3).

#### Threat assessment

Full methodological details and results of this section are described in Goettsch *et al.*[3] Summarizing, during the process 224 taxa were evaluated according to the International Union for Conservation of Nature, IUCN, Red List Categories and Criteria[14] (see Table 1, Supplementary Table 3). The IUCN Red List is a critical indicator to identify species most vulnerable to extinction considering a set of criteria, i.e. species' population trends, size, structure, and geographic ranges. A Red List workshop with the participation of 25 experts from different project partner institutions and IUCN specialists was organized to assess the extinction risk of taxa.

#### Species distribution modeling

To compile occurrence records, hundreds of sources were consulted, including published and personal databases of the project participants,[eg.15–18] the Agrobiodiversity Atlas of Guatemala (https://www.scidev.net/america-latina/biodiversidad/noticias/atlas-digital-cataloga-plantas-silvestres-de-guatemala.html), the Global Biodiversity Information Facility (GBIF, https://www.gbif.org/), and Mexico's Biodiversity Information System (SNIB, http://snib.mx/).

To generate potential species distribution models (SDM), we used more than 18,000 occurrence records (Supplementary Table 4), that were standardized and curated by experts to generate the range maps of taxa as part of the extinction risk assessment, which were published in IUCN Red List.[3] Spatial resolution of the SDM was 1 $km^2$. SDM were obtained for taxa with more than 20 unique occurrence data in a 1 $km^2$ grid covering the study extent in order to reduce uncertainty when using smaller sample sizes.[19] We used bioclimatic variables, such as annual potential evapotranspiration, aridity index, annual radiation, slope and altitude.[20–22] Climate data represents annual and seasonal patterns of climate between 1950 and 2000. Also, we used a variable that described the percentage of bare soil and cultivated areas.[23] Collinearity between variables was assessed with the 'corselect' function of the package fuzzySim version 1.0[24], using a value of 0.8 as threshold to exclude highly correlated variables.

We used MaxEnt, a machine-learning algorithm that uses the maximum entropy principle to identify a target probability distribution, subject to a set of constraints related to the occurrence records and environmental data.[25,26] Model calibration area for each taxon included those ecoregions where the taxon has been recorded; we used the terrestrial ecoregions dataset[6]. We did this based on the calibration area or 'M element' of the BAM diagram that refers to areas that have been accessible to the taxon via dispersal over relevant periods of time.[27,28] We randomly sampled 10,000 background localities from the select areas.

In order to reduce model complexity without compromising performance, we built several models by varying the feature classes (FC) and regularization multipliers (RM) (see[29–31]) using R 3.6.0[32] and 'ENMeval' package.[33] FC determines the flexibility of the modeled response to the predictor variables, while the RM penalizes model complexity.[29] Occurrence records were randomly divided into 70% for model selection, and 30% of data was withheld for model validation. ENMeval carries out an internal partition of localities to test each combination of settings. Therefore, we selected the random k-fold method to divide localities into four bins. We build models with six FC combinations and varied RM values ranging from 0.5 to 4.0 in 0.5 increments. Optimal models were selected using Akaike's Information Criterion corrected for small sample sizes ($\Delta$AICc = 0). This method penalizes overly complex models and helps to choose those with an optimal number of parameters. However, it has been shown that the number of model parameters may not correctly estimate degrees of freedom,[34] and that model selection should not be selected solely with one measure.[35] Thus, we used 30% of the withheld data to test the area under curve (AUC) of the receiver operating characteristic, and the omission error under a 10 percentile training threshold.

Experts selected the best fitting SDM and indicated possible overestimated areas, which were then eliminated case by case using information of Mexican ecoregions[36] and watersheds[37]. For *Phaseolus and Zea,* we used SDM that were previously generated by Delgado-Salinas et al.,[38] and Sánchez González et al.,[39] respectively. Final SDM were validated by experts of each taxonomic group. For analysis purposes, we used 116 SDM (see Supplementary Table 3) and trimmed continuous SDM using binary SDM to keep pixel values of areas with elevated probability of taxa presence and clipped models to the Mexican territory.

#### Proxies of genetic diversity

To capture and spatially delimit potential genetically distinct populations within the geographic range of taxa in order to guide conservation planning of genetic diversity of CWR, we developed a new approach integrating environmental and phylogeographic information. Phylogeographic studies inform on the drivers of intraspecific diversity in several species that assume correlation and interaction between genetics and ecological variables.[eg.40] The information can be used to infer proxies of intraspecific variation and to generate spatial input that is key for prioritization analysis of CWR, as it may be unfeasible to sample or generate genetic analyses of hundreds of taxa due to limited timeframes and conservation budgets.

We used two sets of criteria: a) environmental variability according to Holdridge life zones[41], and b) historic differentiation, as shown by phylogeographic patterns found in other species of the same habitat and region. Studies in Mexico indicate that altitudinal changes as given by mountain ranges are the main drivers affecting climate fluctuations rather than latitudinal changes.[10] In addition, there is strong evidence that genetic structure in species or taxonomic groups can be different in relatively close areas; eg., hummingbird populations distributing east or west of Istmo de Tehuantepec in south Mexico.[40]

For environmental variability, we used Holdridge life zones to characterize climatically distinct areas of Mesoamerica based on biotemperature, annual precipitation and potential evapotranspiration ratio (Supplementary Table 5). To gather data of phylogeographical boundaries that shaped genetic structure in Mexico, we performed a literature review and compiled a list of spatially explicit phylogeographic patterns reported for the country (Supplementary Table 6). To represent phylogeographic patterns not reflected by life zones, we split the life zones into different subzones using best fitting cartography to represent the landscape context, i.e. biogeographic provinces[42], edaphology[43] and watersheds[37] (see Fig. 1). We considered that in the context of CWR conservation planning it is safer to assume genetic variation than to omit heterogeneity in the distribution ranges of taxa. For analysis purposes, we subdivided each SDM by PGD, thus delimiting areas of potential population differentiation, using ArcGIS.[44] Combining 116 SDM with 102 PGD resulted in 11,832 layers; but for further analysis, we used 5,004 input layers with value data, which were filtered by using R 4.0.0.[45]

We validated our findings by using genomic data of an empirical study of maize wild relatives, the teosinte *Zea mays* ssp. *parviglumis.* The dataset includes ca. 1,800 occurrence records and ca. 30,000 SNPs.[46] Sampling localities were not used for distribution modelling. Admixture groups per population were estimated for K1 to 60. We chose K=13 for plotting based on the CV error. The proportion of each genetic clusters was estimated for sampling locality and plotted using pie charts over the map. Then, using a raster of the SDM of the species subdivided by PGD, we extracted which was the PGD most frequent in a 5 km buffer for each sampling locality. The Admixture plot was ordered by all genetic clusters and subdivided by the PGD most frequent for each locality.

#### Habitat preference

We considered habitat preference to refine the presence of CWR in the SCP process; thus minimizing commission errors and highlighting areas that more probably contain taxa.[47] For each taxon, experts assessed its habitat preference (1: high preference; 0.5: low preference; 0.1: no preference) according to the following categories: i) well conserved vegetation (i.e. primary vegetation), ii) human impacted vegetation (i.e. secondary vegetation), iii) less intensive rainfed and moisture agriculture, iv) intensive rainfed and moisture agriculture, v) irrigated agriculture, vi) induced and cultivated grasslands and forests, and vii) urban areas (Supplementary Table 7). To spatially delimit these classes, we used the land use cover and vegetation map for Mexico,[48] and assessed seven main categories of land cover. To differentiate between less intensive and intensive cultivated areas, we followed Bellon et al.[49] who associated the presence of native maize varieties of Mexico to occur in municipalities with average yields of less than or equal to 3 t ha-1. We also used agricultural production data from 2010 from the Information System of Agrifood and Fisheries (SIAP), and selected the municipalities with the established average maize yield. We combined the municipality layer with the land cover map to differentiate areas of high and low agricultural intensity. To generate taxon-specific habitat layers, we associated the habitat preference classes established by experts to the land cover map aggregated into seven major land cover categories, using R 3.6.0[32] and the ‘raster’[50] and ‘rgdal’.[51]

#### Spatial conservation planning analysis

We identified areas of high conservation value for CWR in Mexico by using the software Zonation, a systematic conservation planning tool that allows optimizing representation of species or taxa and other conservation features, e.g. PGD, in a given study area.[52] The program hierarchically ranks areas by removing cells of low conservation value, as given for example by reduced species richness or occurrence of low weighted features, while considering multiple criteria such as habitat preference of taxa. We applied the core-area zonation removal rule (CAZ) to maximize representation of all conservation elements, i.e. weighted taxa as well as range restricted and wide distributing taxa, in a minimal possible area[53] (see Zonation configuration in Supplementary Information 1). Zonation generates two main outputs: (a) a hierarchical map of continuous values, that allows decision makers establishing different area thresholds to highlight areas of conservation interest; and (b) a species' representation curve that shows species or feature range distribution in a given area. The curve also allows identifying how much area is needed to cover a certain area of species' distribution or the distribution of a feature of conservation interest.

We generated five preliminary approaches to assess the input data that maximizes the representation of genetic diversity of CWR, considering: i) 116 trimmed SDM validated by experts of each taxonomic group, which we used as a reference to examine the representation of taxa and proxies of genetic variability; ii) 116 SDM and 27 layers representing Holdridge life zones; iii) 116 SDM and 102 layers representing the PGD individually; iv) 5,004 input layers representing the intersection of SDM and PGD; v) 116 SDM as the main conservation features, while integrating one single layer of PGD in order to assess them as administrative units by using Zonation's Administrative units function; using a strong local representation forces taxon representation within in each unit, here PGD, thus maximizing representation of genetic diversity of each taxon. We performed statistical analysis to compare the results of these five scenarios by establishing area thresholds of 20% of Mexico's terrestrial area, and assessed the scenario with highest feature representation. Statistics were done using R 3.6.0[32] and the packages: 'dplyr'[54], 'ggplot2'[55], 'raster'[50], 'scales'[56], 'sp'[57,58], 'tidyr'[59], and 'vegan'.[60]

For final conservation analyses, we used the following inputs: 1) 5,004 layers, i.e. SDM combined with PGD to maximize representation of genetic diversity, 2) occurrence records of 103 taxa; only for those taxa without SDM (see Supplementary Table 3), 3) taxa specific habitat layers, and 4) IUCN threat category as an additional parameter for taxa (see taxa IUCN threat category at Supplementary Table 3). See Zonation configuration at the end of this document.

We included occurrence data of taxa without SDM to prevent missing important areas of taxa known distribution that are important to conserve. We enabled the function 'species of special interest' (SSI) to include a list of taxa with georeferenced data. The spatial reference system was World Mercator projection.

We assigned weights to taxa with SDM by using IUCN threat categories, giving highest values to taxa with highest risk of extinction that urgently need management actions to further avoid genetic erosion. Thus, weights were assigned as follows: Critically endangered, CR: 1; Endangered, EN: 1; Vulnerable, VU: 0.8; Near threatened, NT: 0.5; Data deficient, DD: 0.3; Least concern, LC: 0.2 Not evaluated, NE: 0.1. As there is no rule for weight setting, we assigned values between 0 and 1 regardless of taxa distribution ranges, which is automatically considered in the Zonation algorithm. SSI taxa were all weighted similarly with 1.

We generated three final scenarios to identify conservation areas for (a) all taxa, (b) taxa exclusively distributing in natural vegetation, and (c) taxa associated to a wider range of habitats such as natural vegetation, agricultural and urban areas. The

Zonation configuration remained similar among the three scenarios. When taxa were not included in a given scenario, we assigned a value of 0, this excluded the feature to be considered for the hierarchical prioritization of the landscape, but still allowed to evaluate the taxa during post-processing.

We discussed the proposed methodological framework, input layer and criteria during a fourth workshop in Mexico. It is worth mentioning that we run several preliminary analyses, which included additional layers, such as indigenous areas that promote the presence of CWR in the landscape.[61] However, as the output indicated no evident difference by including this information, final analyses did not consider these data. We neither included protected areas (PAs) nor tried to expand on the current 12% PA system, because most PA management plans do not specifically address CWR management (but see the PA management program of ‘Sierra de Manantlán’[62]), and thus generally do not adequately plan for wild and native genetic resources.[63]

#### Results

##### CWR inventory

As detailed in Goettsch et al.,[3] the selected Mesoamerican CWR inventory included 224 taxa of nine crops, including chili pepper (four *Capsicum* spp*.)*, squash (11 *Cucurbita* spp.), cotton taxa (13 *Gossypium* spp.), avocado (18 *Persea* spp.), bean (55 *Phaseolus* spp.), husk tomato (67 *Physalis* spp.), potato (26 *Solanum* sect. *Petota.*), 22 maize taxa (11 taxa of *Zea* spp. and 11 taxa of *Tripsacum* spp.), and vanilla (8 *Vanilla* spp.) (see Supplementary Table 3).

##### Threat assessment

Also detailed in Goettsch et al.,[3] the threat analysis included not only species, but subspecies and subpopulations (i.e. races) for some groups (Supplementary Table 3). Using the IUCN Red List Categories and Criteria, one-third (71 spp.) of the evaluated CWR taxa are threatened (CR: 7 spp., EN: 48 spp., and VU: 16 spp.). Twenty taxa were assessed as DD as there was insufficient data to evaluate them. Cotton and vanilla were the most threatened groups with 92% and 89% of their evaluated species, respectively, at risk of extinction.

Threats affecting CWR include habitat loss and degradation -mainly due to the expansion of crops, livestock and infrastructure, such as urban development and roads, native pests, exotic invasive species and genetically modified organisms. Other main threat factors are climate change and extreme weather events. While the impact of these factors may be independent, in general, there are several stressors affecting (agro-)biodiversity, interacting in synergy and in complex ways, thus exacerbating their effects.[3]

##### Taxa distribution patterns

Based on the distribution of occurrence data in a 5 $km^2$ grid, areas of highest number of taxa are located in the central part of Mexico, particularly in the valley of Mexico, in the surroundings of Jalapa and in the region of Los Tuxtlas in the state of Veracruz, in the basin of Tehuacán-Cuicatlán, as well as in the surroundings of San Cristóbal de las Casas, Chiapas, in South Mexico (Supplementary Fig. 1). Unsurprisingly, areas of high taxa richness are related to areas with highest sampling effort. The presence of research centers or biological stations explains the general diversity pattern of CWR based on occurrence records.

For 116 taxa (species and subspecies), SDM were obtained; all were validated by experts (see Supplementary Table 3). See MaxEnt performance and significance of SDM at Supplementary Table 8. AUC values ranged from 0 to 1; 0.5 indicated a model performance not better than random, while values closer to 1 indicated a better model performance; here we used SDM showing AUC values higher than 0.7. Direct download links are available at Supplementary Table 9. Combined binary maps, i.e. presence – absence maps, highlighted areas with the highest number of CWR located in the Trans-Mexican Volcanic Belt, the region of 'Los Altos de Chiapas', and the Soconusco region, in southern Mexico (Fig. 1).

Considering the occurrence data and SDM of taxa within the context of protected areas (PAs), our results show that potentially there are seven PAs with more than 20 taxa (Supplementary Table 10). The PAs with highest numbers of recorded taxa are C.A.D.N.R. 043 Estado de Nayarit (46 taxa), and Sierra de Manantlán (42 taxa); they also represent the PAs with highest numbers of taxa based on SDM. However, most of the PAs had considerable differences between the observed and estimated number of taxa, see for example Sierra de Quila (4 vs. 65 taxa, respectively), Insurgente José María Morelos (5 vs. 62 taxa, respectively), and Sierra de Huautla (6 vs. 60 taxa, respectively). Only nine PAs indicate that approximately half of the estimated number of taxa have actually been recorded, including the PAs of Revillagigedo, Bahía de Loreto, and El Pinacate y Gran Desierto de Altar. In these areas on average less than five species have been reported, but up to 11 taxa are potentially distributed in these PAs. SDM represent sites with environmental conditions similar to where the species has been observed, i.e. presence data rather than absence information is used to obtain SDM, thus estimating the commission error is difficult.[64]

#### Proxies of genetic diversity

We established 102 PGD in Mexico based on environmental variability and historic differentiation (Supplementary Fig. 3). The PGD are characterized by specific ranges of temperature, humidity and potential evapotranspiration according to Holdridge life zones characterization (Supplementary Table 5), and reveal genetic differentiation based on phylogeographic patterns (Supplementary Table 6). To test our approach, we used distribution data of *Zea mays* ssp. *parviglumis* and assessed the performance of proxies. The distribution of maize mainly occurs in 12 PGD; the number rises up to 28 proxies if isolated pixels of other proxies are taken into account (Supplementary Fig. 4). In comparison, the analysis of genetic diversity along the geographical range of the taxon revealed 13 populations (Sánchez per. com.). While there was no full coincidence of estimated versus observed number of populations, the use of PGD allows illustrating the genetic variation within the distribution ranges of taxa (Fig. 3).

#### Conservation areas

Using SDM combined with PGD showed the highest representation of genetic diversity (Fig. 4). Other evaluated approaches were not as efficient to represent genetic variation within taxa ranges, only employing SDM in the analysis showed least representation of PGD as given by mean proportion of area of proxies represented by taxa distribution. Therefore, we used the combined input of SDM and PGD, i.e. 5,004 raster layers.

Areas for conservation of Mesoamerican CWR are located throughout the entire country, occurring in different environmental and socio-economic conditions (Supplementary Fig. 6). Although distribution patterns of the conservation areas differ when considering (a) all taxa, (b) taxa exclusively distributed in natural vegetation, and (c) taxa associated to different habitats, there are still common areas across the three scenarios (Supplementary Fig. 7). Large aggregated areas are located in the temperate mountain ranges of the Trans-Mexican Volcanic Belt, characterized by high taxon richness; in the region ranging from central Veracruz state to Chiapas, crossing Puebla and including large areas of Oaxaca, the Tehuacán-Cuicatlán valley and the Chimalapas region, corresponding to areas of heterogeneous environments; and along the northern coastline of the Michoacán state in Central Mexico, and in the cloud forests and humid tropical areas of southern Mexico, representing habitats for species such as *Vanilla odorata*.

When considering the three scenarios designed for Mesoamerican CWR (i.e. (a) all taxa, (b) taxa exclusively distributed in natural vegetation, and (c) taxa associated to different habitats, there are still common areas across the three scenarios), representation curves of distribution ranges of taxa revealed that a significant portion of CWR ranges can be represented in a fraction of Mexico's terrestrial area (Supplementary Fig. 8). Average values differ between groups due to the conservation weight. Although taxa with high risk of extinction (CR and EN) had highest conservation weights, which generally result in highest representation, here the results show a different pattern as their potential distribution ranges are highly impacted by anthropogenic factors, such as forest loss or degradation. Consequently, it was not possible to represent the 100% of the potential distribution of taxa. Performance curves quantifying the proportion of priority taxa within the scenario considering all priority taxa are shown in Supplementary Fig. 9 (see scenario in Supplementary Fig. 6a)

Considering the three different scenarios, the results expose habitat preferences of taxa targeted in each scenario. The scenario considering taxa exclusively distributed in natural vegetation showed a higher proportion of area in primary and secondary vegetation, while the scenario based on taxa associated to different habitats showed a higher proportion of area in rainfed and moisture agriculture. This is evident when analyzing 20% of the area of each scenario (Supplementary Fig. 10).

Also, within 20% of Mexico's area, we found that almost half of the area is located inside indigenous areas, and 11% of the selected area is covered by federal protected areas (Supplementary Fig. 11; Supplementary Tables 10 and 11).

Notably, all conservation features, i.e. taxa's distribution range as given by PGD, are represented in the selected 20% of area; on average, 50% of their spatial extent. This means that the solutions are effective to represent the genetic diversity of a given crop in the top fraction of land. Nonetheless, to represent at least 50% of all conservation features, 80% of Mexico's terrestrial surface must be sustainably managed, independently of the scenario.

#### Zonation configuration

##### Settings file

[Settings]
removal rule = 1 (# Core Area Zonation)
warp factor  = 1
edge removal = 1
annotate name = 0

use SSI = 1
SSI file name = SSI_list.txt

use groups = 1
groups file = habitat_group_features.txt

use condition layer = 1
condition file = habitat_features.txt

##### Batch file (run in cluster)

#!/bin/bash
zig4 -r E_final.dat features_Todos.spp /home/see/E_final/E_final_Todos.txt 0.0 0 0 0 (# analysis for all taxa)

#!/bin/bash
zig4 -r E_final.dat features_VegPyS.spp /home/see/E_final/E_final_VegPyS.txt 0.0 0 0 0  (# analysis for taxa exclusively distributed in well-preserved vegetation)

#!/bin/bash
zig4 -r E_final.dat features_HabVarios.spp /home/see/E_final/E_final_HabVarios.txt 0.0 0 0 0 (# analysis for taxa that can be associated to different habitats and land uses (e.g. natural vegetation, agriculture and urban areas)
