## Supplementary Materials 2 for "Incorporating evolutionary and threat processes into crop wild relatives conservation"

### Supplementary Information 2

#### Supplementary Tables and Figures

##### Index of supplementary tables

Supplementary Table 1.CWR checklist. (provided as separate file)

Supplementary Table 2. CWR subset. (provided as separate file)

Supplementary Table 3. Inventory of selected Mesoamerican CWR, IUCN Red List Criteria and Category (see details in Goettsch *et al.* in prep.), and spatial format used for analysis (SDM: species distribution model; Occs: occurrence georeferenced data). (IUCN Red List Category, CR: Critically endangered, EN: Endangered, VU: Vulnerable, NT: Near threatened, LC: Least concern, DD: Data deficiency).

Supplementary Table 4. CWR occurrence records. (provided as separate file)

Supplementary Table 5. Holdridge life zone characterization of Mexico.

Supplementary Table 6. Selected references of phylogeographic patterns reported for Mexico used to assess proxies of genetic diversity.

Supplementary Table 7. Habitat preferences of Mesoamerican CWR with potential species distribution model used in the spatial analysis (1: high preference; 0.5: low preference; 0.1: no preference).

Supplementary Table 8. MaxEnt performance and significance of potential species distribution models (SDM). (Occurrence data was divided into 70% for training and 30% for testing the model. MaxEnt performance was tested with 30% of the withheld data. Testing omission rate is given for SDM selected by experts, which were mainly model threshold with the ten percentile training presence.) (*indicates omission rates under a minimum training presence threshold.)

Supplementary Table 9. Direct download links for CWR potential species distribution models.

Supplementary Table 10. CWR in protected areas. Information is based on occurrences (occs.) of 224 taxa and 116 potential species distribution models (SDM).

Supplementary Table 11. Area of three scenarios for conservation of Mesoamerican CWR in Mexico in federal protected areas and indigenous areas. For analysis, 20% of Mexico's terrestrial area is considered of each scenario.

**Index of supplementary figures**

Supplementary Fig. 1. Spatial pattern of taxa richness of selected Mesoamerican CWR in Mexico based on occurrence georeferenced data. Spatial resolution $5km^2$.

Supplementary Fig. 2. Holdridge life zones of Mexico, based on biotemperature, annual precipitation and potential evapotranspiration ratio (see Supplementary Table 5). Life zones are represented by different colours (see code of numbers at Supplementary Table 5). Spatial resolution $1km^2$.

Supplementary Fig. 3. Proxies of genetic diversity (PGD) for Mexico, based on environmental data (Supplementary Table 5, Supplementary Fig. 2) and historic drivers (Supplementary Table 6). The 102 PGD are represented by different colours. Spatial resolution $1km^2$.

Supplementary Fig. 4. Potential species distribution model of *Z. mays* ssp. *parviglumis* as given by PGD (background colours).

Supplementary Fig. 5. Land cover map that was used to assess habitat preference of each taxon (Supplementary Table 7). Spatial resolution $1km^2$.

Supplementary Fig. 6. Conservation areas for Mesoamerican crop wild relatives in Mexico, considering (a) all taxa, (b) taxa exclusively distributing in natural vegetation, and (c) taxa associated to different habitats. Spatial resolution $1km^2$.

Supplementary Fig. 7. Coincidence of three conservation scenarios, considering (a) all taxa; (b) taxa exclusively distributing in natural vegetation; and (c) taxa associated to different habitats. Twenty percent of Mexico's terrestrial area is highlighted of each scenario (see continuous values at Supplementary Fig. 6). Spatial resolution $1km^2$.

Supplementary Fig. 8. Performance curves quantifying the proportion of taxa distribution ranges considered for each scenario, grouped by IUCN Red List Category. Scenarios considered (a) all taxa; (b) taxa exclusively distributing in natural vegetation; and (c) taxa associated to different habitats, i.e. natural vegetation, agricultural and urban areas.

Supplementary Fig. 9. Performance curves quantifying the proportion of taxa distribution ranges considering all priority taxa (see Fig. 5a or Supplementary Fig. 6a), grouped by genus; colours are according to its IUCN Red List Category.

Supplementary Fig. 10. Conservation areas for Mesoamerican crop wild relatives in Mexico according to land cover data used for the analysis (see Supplementary Fig. 5). Scenarios are based on proxies of genetic diversity, and consider (a) all taxa, (b) taxa exclusively distributing in natural vegetation, and (c) taxa associated to different habitats. Twenty percent of Mexico's terrestrial area is highlighted of each scenario. Spatial resolution $1km^2$.

Supplementary Fig. 11. Conservation areas for Mesoamerican crop wild relatives in Mexico, considering (a) all taxa, (b) taxa exclusively distributing in natural vegetation, and (c) taxa associated to different habitats, showing continuous values (left), and highlighting 20% of Mexico's terrestrial area, federal protected areas and indigenous areas. Spatial resolution $1km^2$.

**Supplementary Table 3. Inventory of selected Mesoamerican CWR, IUCN Red List Criteria and Category (see details in Goettsch *et al.* in prep.), and spatial format used for analysis purposes (SDM: species distribution model; Occs: occurrence georeferenced data). (IUCN Red List Category, CR: Critically endangered, EN: Endangered, VU: Vulnerable, NT: Near threatened, LC: Least concern, DD: Data deficiency).**

| **Taxon** | **IUCN Red List Criteria** | **IUCN Red List Category** | **SDM / Occs.** |
|---|---|---|---|
| *Capsicum annuum* | N/A | LC | SDM |
| *Capsicum frutescens* | N/A | LC | SDM |
| *Capsicum lanceolatum* | B2ab(iii,iv,v) | EN | SDM |
| *Capsicum rhomboideum* | N/A | LC | SDM |
| *Cucurbita argyrosperma* | N/A | LC | SDM |
| *Cucurbita cordata* | N/A | DD | SDM |
| *Cucurbita digitata* | N/A | LC | SDM |
| *Cucurbita foetidissima* | N/A | LC | SDM |
| *Cucurbita lundelliana* | N/A | LC | SDM |
| *Cucurbita okeechobeensis* subsp. *martinezii* | N/A | NT | SDM |
| *Cucurbita palmata* | N/A | DD | SDM |
| *Cucurbita pedatifolia* | N/A | DD | SDM |
| *Cucurbita pepo* | N/A | LC | |
| *Cucurbita pepo* subsp. *fraterna* | B1a | NT | SDM |
| *Cucurbita radicans* | B2ab(iii) | EN | SDM |
| *Gossypium aridum* | B2ab(i,ii,iii,iv,v) | VU | SDM |
| *Gossypium armourianum* | C2a(i) | CR | Occs. |
| *Gossypium barbadense* | N/A | LC | Occs. |
| *Gossypium davidsonii* | B2ab(i,ii,iii,v) | VU | SDM |
| *Gossypium gossypioides* | B1ab(iii,v); C2a(i) | VU | SDM |
| *Gossypium harknessii* | B2ab(i,ii,iii,iv,v) | EN | SDM |
| *Gossypium hirsutum* | B2ab(i,iii,v); C1+2a(i) | VU | SDM |
| *Gossypium laxum* | B1ab(iii,v)+2ab(iii,v); C2a(i); D | EN | Occs. |
| *Gossypium lobatum* | A4ac; B1ab(i,ii,iii,v)+2ab(i,ii,iii,v); C2a(i) | EN | Occs. |
| *Gossypium schwendimanii* | A4ac; B1ab(i,ii,iii,v)+2ab(i,ii,iii,v); C2a(i) | EN | Occs. |
| *Gossypium thurberi* | B2ab(i,ii,iii) | EN | SDM |
| *Gossypium trilobum* | B1ab(ii,iii,v)+2ab(ii,iii,v); D | EN | Occs. |
| *Gossypium turneri* | B1ab(i,ii,iii,v); C2a(i) | CR | Occs. |
| *Persea albida* | A2c | EN | SDM |

| | | | |
|---|---|---|---|
| *Persea americana* | N/A | LC | SDM |
| *Persea caerulea* | N/A | LC | SDM |
| *Persea chamissonis* | A2c | EN | Occs. |
| *Persea cinerascens* | A2c; B2ab(i,ii,iii,v) | EN | Occs. |
| *Persea donnell-smithii* | A2c | VU | SDM |
| *Persea hintonii* | A2c | VU | SDM |
| *Persea liebmannii* | N/A | LC | SDM |
| *Persea longipes* | A2c; B2ab(iii) | EN | SDM |
| *Persea pallescens* | A2c; B2ab(iii) | EN | SDM |
| *Persea podadenia* | A2c | NT | SDM |
| *Persea purpusii* | N/A | DD | SDM |
| *Persea rigens* | N/A | LC | |
| *Persea rufescens* | A2c; B2ab(iii) | EN | Occs. |
| *Persea schiedeana* | A2c | EN | SDM |
| *Persea sessilis* | N/A | DD | |
| *Persea veraguasensis* | N/A | DD | |
| *Persea vesticula* | N/A | LC | SDM |
| *Phaseolus acutifolius* | N/A | LC | SDM for two subspecies |
| *Phaseolus albescens* | B1ab(i,iii,v) | VU | Occs. |
| *Phaseolus albiflorus* | B1ab(iii,v) | VU | Occs. |
| *Phaseolus amblyosepalus* | N/A | LC | Occs. |
| *Phaseolus angustissimus* | N/A | LC | SDM |
| *Phaseolus anisophyllus* | N/A | DD | Occs. |
| *Phaseolus campanulatus* | D | EN | Occs. |
| *Phaseolus carterae* | B1ab(iii) | EN | Occs. |
| *Phaseolus chiapasanus* | D | EN | SDM |
| *Phaseolus coccineus* | N/A | LC | SDM |
| *Phaseolus dasycarpus* | B1ab(iii,v) | EN | Occs. |
| *Phaseolus dumosus* | B1ab(iii,v) | EN | Occs. |
| *Phaseolus esperanzae* | D1 | VU | Occs. |
| *Phaseolus filiformis* | N/A | LC | SDM |
| *Phaseolus glabellus* | N/A | LC | SDM |
| *Phaseolus hintonii* | D | EN | Occs. |
| *Phaseolus jaliscanus* | N/A | LC | SDM |
| *Phaseolus juquilensis* | N/A | LC | Occs. |
| *Phaseolus laxiflorus* | N/A | LC | Occs. |
| *Phaseolus leptophyllus* | N/A | DD | Occs. |
| *Phaseolus leptostachyus* | N/A | LC | SDM |
| *Phaseolus lunatus* | N/A | LC | SDM |
| *Phaseolus macrolepis* | B1ab(iii,v) | EN | |
| *Phaseolus maculatifolius* | N/A | LC | Occs. |
| *Phaseolus maculatus* | N/A | LC | SDM |
| *Phaseolus macvaughii* | N/A | LC | Occs. |
| *Phaseolus marechalii* | N/A | LC | Occs. |

| | | | |
|---|---|---|---|
| *Phaseolus micranthus* | N/A | LC | SDM |
| *Phaseolus microcarpus* | N/A | LC | SDM |
| *Phaseolus neglectus* | N/A | LC | Occs. |
| *Phaseolus nelsonii* | D1 | VU | Occs. |
| *Phaseolus nodosus* | N/A | LC | Occs. |
| *Phaseolus novoleonensis* | B1ab(iii) | EN | Occs. |
| *Phaseolus oaxacanus* | N/A | LC | Occs. |
| *Phaseolus oligospermus* | D1 | VU | Occs. |
| *Phaseolus parvifolius* | N/A | LC | SDM |
| *Phaseolus parvulus* | N/A | LC | SDM |
| *Phaseolus pauciflorus* | N/A | LC | SDM |
| *Phaseolus pedicellatus* | N/A | LC | SDM |
| *Phaseolus perplexus* | N/A | LC | SDM |
| *Phaseolus plagiocylix* | B1ab(i,iii,v) | EN | Occs. |
| *Phaseolus pluriflorus* | N/A | LC | SDM |
| *Phaseolus purpusii* | B2ab(iii,v) | EN | Occs. |
| *Phaseolus reticulatus* | B1ab(iii)+2ab(iii) | EN | Occs. |
| *Phaseolus ritensis* | N/A | LC | SDM |
| *Phaseolus rotundatus* | N/A | LC | Occs. |
| *Phaseolus salicifolius* | N/A | LC | Occs. |
| *Phaseolus scabrellus* | N/A | LC | SDM |
| *Phaseolus sonorensis* | N/A | LC | Occs. |
| *Phaseolus tenellus* | D | EN | Occs. |
| *Phaseolus tuerckheimii* | N/A | LC | Occs. |
| *Phaseolus vulgaris* | N/A | LC | SDM |
| *Phaseolus xanthotrichus* | N/A | LC | Occs. |
| *Phaseolus xolocotzii* | N/A | LC | Occs. |
| *Phaseolus zimapanensis* | N/A | LC | Occs. |
| *Physalis acutifolia* | N/A | LC | SDM |
| *Physalis aggregata* | B1ab(iii) | VU | Occs. |
| *Physalis ampla* | N/A | LC | Occs. |
| *Physalis angulata* | N/A | LC | SDM |
| *Physalis angustior* | N/A | DD | Occs. |
| *Physalis angustiphysa* | N/A | LC | SDM |
| *Physalis campanula* | B1ab(ii,iii) | NT | Occs. |
| *Physalis campechiana* | N/A | LC | SDM |
| *Physalis caudella* | N/A | LC | SDM |
| *Physalis chenopodiifolia* | N/A | LC | SDM |
| *Physalis cinerascens* | N/A | LC | SDM |
| *Physalis cinerascens* var. *cinerascens* | N/A | LC | |
| *Physalis cinerascens* var. *spathulifolia* | N/A | LC | |
| *Physalis cordata* | N/A | LC | SDM |
| *Physalis coztomatl* | N/A | LC | SDM |

| | | | |
|---|---|---|---|
| *Physalis crassifolia* | N/A | LC | SDM |
| *Physalis crassifolia* var. *infundibularis* | N/A | LC | |
| *Physalis crassifolia* var. *versicolor* | N/A | LC | |
| *Physalis glabra* | N/A | LC | Occs. |
| *Physalis glutinosa* | N/A | LC | SDM |
| *Physalis gracilis* | N/A | LC | SDM |
| *Physalis greenmanii* | B1ab(iii) | EN | Occs. |
| *Physalis hastatula* | B1ab(iii) | EN | Occs. |
| *Physalis hederifolia* | N/A | LC | SDM |
| *Physalis hintonii* | N/A | LC | Occs. |
| *Physalis hunzikeriana* | N/A | DD | Occs. |
| *Physalis ignota* | N/A | LC | SDM |
| *Physalis lagascae* | N/A | LC | SDM |
| *Physalis lassa* | N/A | LC | Occs. |
| *Physalis latecorollata* | | DD | Occs. |
| *Physalis latiphysa* | N/A | LC | Occs. |
| *Physalis leptophylla* | N/A | LC | SDM |
| *Physalis lignescens* | B1ab(iii) | EN | Occs. |
| *Physalis lobata* | N/A | LC | Occs. |
| *Physalis longicaulis* | | DD | Occs. |
| *Physalis longiloba* | N/A | LC | Occs. |
| *Physalis longipedicellata* | N/A | LC | Occs. |
| *Physalis mcvaughii* | N/A | NT | Occs. |
| *Physalis melanocystis* | N/A | LC | SDM |
| *Physalis microcarpa* | N/A | LC | Occs. |
| *Physalis microphysa* | N/A | LC | Occs. |
| *Physalis minimaculata* | B1ab(iii) | VU | Occs. |
| *Physalis minuta* | N/A | LC | SDM |
| *Physalis muelleri* | B2ab(iii) | CR | Occs. |
| *Physalis nicandroides* | N/A | LC | SDM |
| *Physalis orizabae* | N/A | LC | SDM |
| *Physalis parvianthera* | N/A | DD | Occs. |
| *Physalis patula* | N/A | LC | SDM |
| *Physalis pennellii* | B1a | NT | Occs. |
| *Physalis philadelphica* | N/A | LC | SDM |
| *Physalis philippensis* | N/A | DD | Occs. |
| *Physalis porrecta* | N/A | DD | Occs. |
| *Physalis pringlei* | N/A | LC | Occs. |
| *Physalis pruinosa* | N/A | LC | SDM |
| *Physalis pubescens* | N/A | LC | SDM |
| *Physalis queretaroensis* | N/A | LC | Occs. |
| *Physalis rydbergii* | N/A | DD | Occs. |
| *Physalis sancti-josephi* | N/A | DD | Occs. |
| *Physalis solanacea* | N/A | LC | SDM |

| | | | |
|---|---|---|---|
| *Physalis sordida* | N/A | LC | SDM |
| *Physalis subrepens* | N/A | LC | Occs. |
| *Physalis sulphurea* | N/A | LC | SDM |
| *Physalis tamayoi* | B1a | NT | Occs. |
| *Physalis tehuacanensis* | B2ab(iii) | CR | Occs. |
| *Physalis virginiana* | N/A | LC | SDM |
| *Physalis volubilis* | N/A | LC | SDM |
| *Physalis waterfallii* | N/A | LC | SDM |
| *Solanum agrimonifolium* | N/A | LC | SDM |
| *Solanum bulbocastanum* | N/A | LC | SDM |
| *Solanum cardiophyllum* | N/A | LC | SDM |
| *Solanum clarum* | B1ab(iii) | VU | SDM |
| *Solanum demissum* | N/A | LC | SDM |
| *Solanum ehrenbergii* | N/A | LC | SDM |
| *Solanum guerreroense* | N/A | DD | Occs. |
| *Solanum hintonii* | C2a(i) | NT | Occs. |
| *Solanum hjertingii* | N/A | LC | SDM |
| *Solanum hougasii* | N/A | LC | SDM |
| *Solanum iopetalum* | N/A | LC | SDM |
| *Solanum jamesii* | N/A | LC | Occs. |
| *Solanum lesteri* | N/A | DD | Occs. |
| *Solanum michoacanum* | B1ab(iii)+2ab(iii) | EN | Occs. |
| *Solanum morelliforme* | N/A | LC | SDM |
| *Solanum oxycarpum* | B2ab(iii) | EN | SDM |
| *Solanum pinnatisectum* | N/A | LC | SDM |
| *Solanum polyadenium* | N/A | LC | SDM |
| *Solanum sambucinum* | N/A | LC | Occs. |
| *Solanum schenckii* | B2ab(iii) | EN | SDM |
| *Solanum stenophyllidium* | N/A | LC | SDM |
| *Solanum stoloniferum* | N/A | LC | SDM |
| *Solanum tarnii* | B1ab(iii) | EN | Occs. |
| *Solanum trifidum* | B1a | NT | SDM |
| *Solanum vallis-mexici* | B1ab(iii)+2ab(iii) | EN | Occs. |
| *Solanum verrucosum* | N/A | LC | SDM |
| *Tripsacum andersonii* | N/A | LC | Occs. |
| *Tripsacum bravum* | B1ab(iii)+2ab(iii) | VU | Occs. |
| *Tripsacum intermedium* | B2ab(iii) | EN | Occs. |
| *Tripsacum jalapense* | N/A | LC | Occs. |
| *Tripsacum lanceolatum* | N/A | LC | SDM |
| *Tripsacum latifolium* | N/A | LC | SDM |
| *Tripsacum laxum* | N/A | LC | Occs. |
| *Tripsacum maizar* | B2ab(iii,v) | EN | Occs. |
| *Tripsacum manisuroides* | N/A | LC | Occs. |
| *Tripsacum pilosum* | N/A | LC | SDM |

| | | | |
|---|---|---|---|
| *Tripsacum zopilotense* | B1ab(iii)+2ab(iii) | EN | Occs. |
| *Vanilla cribbiana* | A3cd; C2a(i); D | CR | Occs. |
| *Vanilla hartii* | A2cd; B2ab(i,ii,iii,v) | EN | Occs. |
| *Vanilla helleri* | N/A | DD | |
| *Vanilla inodora* | B2ab(iii) | EN | Occs. |
| *Vanilla insignis* | B2ab(iii,v) | EN | Occs. |
| *Vanilla odorata* | B2ab(iii,v) | EN | SDM |
| *Vanilla phaeantha* | B2ab(iii,v) | EN | Occs. |
| *Vanilla planifolia* | B2ab(iii,v) | EN | Occs. |
| *Vanilla pompona* | B2ab(iii,v) | EN | SDM |
| *Zea diploperennis* | B1ab(ii,iii,iv,v)c(iii,iv)+ 2ab(ii,iii,iv,v)c(iii,iv) | EN | SDM |
| *Zea luxurians* | B2ab(i,ii,iii,v) | VU | SDM |
| *Zea mays* | N/A | LC | |
| *Zea mays* subsp. *huehuetenangensis* | B1ab(i,ii,iii,v)+2ab(i,ii, iii,v) | EN | SDM |
| *Zea mays* subsp. *mexicana* | N/A | LC | |
| *Zea mays* subsp. *mexicana* Chalco subpopulation | N/A | LC | SDM |
| *Zea mays* subsp. *mexicana* Durango subpopulation | B1ab(iii,v)+2ab(iii,v) | EN | SDM |
| *Zea mays* subsp. *mexicana* Mesa-Central subpopulation | N/A | LC | SDM |
| *Zea mays* subsp. *mexicana* Nobogame subpopulation | B1ab(iii,v) | CR | SDM |
| *Zea mays* subsp. *parviglumis* | N/A | LC | SDM |
| *Zea perennis* | A2ac | CR | SDM |

**Supplementary Table 5. Holdridge life zones characterization of Mexico.**

| Id | Life zone | Temperature (°C) | Annual precipitation (mm) | Potential evapotranspiration |
|---|---|---|---|---|
| 117 | Polar desert | < 1.5 | < 125 | 0.5- 1.0 |
| 216 | Subpolar dry tundra | 1.5-3.0 | < 125 | 1.0- 2.0 |
| 359 | Boreal rain forest | 3.0-6.0 | 1000-2000 | > 0.25 |
| 436 | Cool temperate steppe | 6.0-12.0 | 250-500 | 1.0- 2.0 |
| 447 | Cool temperate moist forest | 6.0-12.0 | 500-1000 | 0.5- 1.0 |
| 458 | Cool temperate wet forest | 6.0-12.0 | 1000-2000 | 0.25-0.5 |
| 469 | Cool temperate rain forest | 6.0-12.0 | 2000-4000 | > 0.25 |
| 513 | Warm temperate desert | 12.0-18.0 | < 125.00 | 8.0- 16.0 |
| 524 | Warm temperate desert | 12.0-18.0 | 125-250 | 4.0- 8.0 |
| 535 | Warm temperate thorn scrub | 12.0-18.0 | 250-500 | 2.0- 4.0 |
| 546 | Warm temperate dry forest | 12.0-18.0 | 500-1000 | 1.0- 2.0 |
| 557 | Warm temperate moist forest | 12.0-18.0 | 1000-2000 | 0.5- 1.0 |
| 568 | Warm temperate wet forest | 12.0-18.0 | 2000-4000 | 0.25-0.5 |
| 613 | Subtropical desert | 18.0-24.0 | < 125 | 8.0- 16.0 |
| 624 | Subtropical desert scrub | 18.0-24.0 | 125-250 | 4.0- 8.0 |
| 635 | Subtropical thorn woodland | 18.0-24.0 | 250-500 | 2.0- 4.0 |
| 646 | Subtropical dry forest | 18.0-24.0 | 500-1000 | 1.0- 2.0 |
| 657 | Subtropical moist forest | 18.0-24.0 | 1000-2000 | 0.5- 1.0 |
| 668 | Subtropical wet forest | 18.0-24.0 | 2000-4000 | 0.25-0.5 |
| 679 | Subtropical rainforest | 18.0-24.0 | 4000-8000 | > 0.25 |
| 712 | Tropical desert | > 24.0 | < 125 | 16.0- 32.0 |
| 723 | Tropical desert scrub | > 24.0 | 125-250 | 8.0- 16.0 |
| 734 | Tropical thorn woodland | > 24.0 | 250-500 | 4.0- 8.0 |
| 745 | Tropical very dry forest | > 24.0 | 500-000 | 2.0- 4.0 |
| 756 | Tropical dry forest | > 24.0 | 1000-2000 | 1.0- 2.0 |
| 767 | Tropical moist forest | > 24.0 | 2000-4000 | 0.5- 1.0 |
| 778 | Tropical wet forest | > 24.0 | 4000-8000 | 0.25-0.5 |

**Supplementary Table 6. Selected references of phylogeographic patterns reported for Mexico used to assess proxies of genetic diversity.**

**Supplementary Table 7. Habitat preferences of Mesoamerican CWR with potential species distribution model used in the spatial analysis (1: high preference; 0.5: low preference; 0.1: no preference).**

| Taxon | Well conserved vegetation | Human impacted vegetation | Less intensive rainfed and moisture agriculture | Intensive rainfed and moisture agriculture | Irrigated agriculture | Induced and cultivated grasslands and forests | Urban areas |
|---|---|---|---|---|---|---|---|
| *Capsicum annuum* | 1 | 1 | 1 | 0.1 | 0.1 | 0.1 | 1 |
| *Capsicum frutescens* | 1 | 1 | 1 | 0.1 | 0.1 | 0.1 | 1 |
| *Capsicum lanceolatum* | 1 | 0.5 | 0.1 | 0.1 | 0.1 | 0.1 | 0.1 |
| *Capsicum rhomboideum* | 1 | 1 | 0.1 | 0.1 | 0.1 | 0.1 | 0.1 |
| *Cucurbita argyrosperma* | 1 | 1 | 1 | 1 | 0.1 | 1 | 1 |
| *Cucurbita cordata* | 1 | 0.5 | 0.5 | 0.1 | 0.1 | 0.1 | 0.1 |
| *Cucurbita digitata* | 1 | 1 | 0.5 | 0.1 | 0.1 | 0.1 | 1 |
| Cucurbita foetidissima | 1 | 1 | 1 | 1 | 0.1 | 1 | 1 |
| *Cucurbita lundelliana* | 1 | 1 | 1 | 1 | 0.1 | 0.5 | 1 |
| *Cucurbita okeechobeensis* subsp. *martinezii* | 1 | 1 | 1 | 0.5 | 0.1 | 1 | 1 |
| *Cucurbita palmata* | 1 | 0.5 | 0.5 | 0.1 | 0.1 | 0.1 | 0.5 |
| *Cucurbita pedatifolia* | 1 | 1 | 1 | 0.1 | 0.1 | 0.1 | 1 |
| *Cucurbita pepo* subsp. *fraterna* | 1 | 1 | 1 | 0.5 | 0.1 | 0.1 | 0.5 |
| *Cucurbita radicans* | 1 | 1 | 1 | 0.1 | 0.1 | 0.1 | 0.5 |
| *Gossypium aridum* | 1 | 1 | 1 | 0.1 | 0.1 | 1 | 0.1 |
| *Gossypium davidsonii* | 1 | 1 | 1 | 0.1 | 0.1 | 0.1 | 0.1 |
| *Gossypium gossypioides* | 1 | 0.1 | 0.1 | 0.1 | 0.1 | 0.1 | 0.1 |
| *Gossypium harknessii* | 1 | 1 | 0.1 | 0.1 | 0.1 | 0.1 | 0.1 |
| *Gossypium hirsutum* | 1 | 1 | 1 | 1 | 0.1 | 1 | 1 |
| *Gossypium thurberi* | 1 | 1 | 0.1 | 0.1 | 0.1 | 0.1 | 0.1 |
| *Persea albida* | 1 | 0.1 | 0.1 | 0.1 | 0.1 | 0.1 | 0.1 |
| *Persea americana* | 1 | 1 | 0.5 | 0.5 | 0.5 | 0.5 | 0.5 |
| *Persea caerulea* | 1 | 1 | 0.1 | 0.1 | 0.1 | 0.1 | 0.1 |
| *Persea donnell-smithii* | 1 | 1 | 0.1 | 0.1 | 0.1 | 0.1 | 0.1 |
| *Persea hintonii* | 1 | 0.1 | 0.1 | 0.1 | 0.1 | 0.1 | 0.1 |
| *Persea liebmannii* | 1 | 0.1 | 0.1 | 0.1 | 0.1 | 0.1 | 0.1 |
| *Persea longipes* | 1 | 1 | 0.1 | 0.1 | 0.1 | 0.1 | 0.1 |
| *Persea pallescens* | 1 | 0.1 | 0.1 | 0.1 | 0.1 | 0.1 | 0.1 |
| *Persea podadenia* | 1 | 1 | 0.1 | 0.1 | 0.1 | 0.1 | 0.1 |
| *Persea purpusii* | 1 | 0.1 | 0.1 | 0.1 | 0.1 | 0.1 | 0.1 |
| *Persea schiedeana* | 1 | 1 | 1 | 0.1 | 0.1 | 0.1 | 0.1 |
| *Persea vesticula* | 1 | 0.1 | 0.1 | 0.1 | 0.1 | 0.1 | 0.1 |
| *Phaseolus acutifolius tenuifolius* | 0.5 | 0.5 | 0.1 | 0.1 | 0.1 | 1 | 0.1 |
| *Phaseolus albescens* | 0.1 | 0.5 | 0.1 | 0.1 | 0.1 | 0.1 | 0.1 |

| | | | | | | | |
|---|---|---|---|---|---|---|---|
| *Phaseolus angustissimus* | 1 | 1 | 0.1 | 0.1 | 0.1 | 0.1 | 0.1 |
| *Phaseolus chiapasanus* | 1 | 0.5 | 0.5 | 0.1 | 0.1 | 0.1 | 0.5 |
| *Phaseolus coccineus* | 0.5 | 0.5 | 0.5 | 0.5 | 0.1 | 0.5 | 0.5 |
| *Phaseolus filiformis* | 1 | 1 | 0.1 | 0.1 | 0.1 | 0.1 | 0.1 |
| *Phaseolus glabellus* | 1 | 0.1 | 0.1 | 0.1 | 0.1 | 0.1 | 0.5 |
| *Phaseolus jaliscanus* | 1 | 1 | 0.5 | 0.1 | 0.1 | 0.5 | 0.1 |
| *Phaseolus leptostachyus* | 1 | 1 | 0.5 | 0.5 | 0.1 | 0.1 | 0.1 |
| *Phaseolus lunatus* | 1 | 1 | 0.5 | 0.5 | 0.1 | 0.1 | 0.5 |
| *Phaseolus maculatus* | 0.5 | 1 | 0.5 | 0.1 | 0.1 | 0.5 | 0.1 |
| *Phaseolus micranthus* | 1 | 1 | 0.1 | 0.1 | 0.1 | 0.1 | 0.1 |
| *Phaseolus microcarpus* | 1 | 0.5 | 0.1 | 0.1 | 0.1 | 0.1 | 0.5 |
| *Phaseolus parvifolius* | 1 | 1 | 0.1 | 0.1 | 0.1 | 0.1 | 0.1 |
| *Phaseolus parvulus* | 1 | 1 | 0.1 | 0.1 | 0.1 | 0.1 | 0.1 |
| *Phaseolus pauciflorus* | 1 | 0.1 | 0.1 | 0.1 | 0.1 | 0.1 | 0.1 |
| *Phaseolus pedicellatus* | 1 | 1 | 0.1 | 0.1 | 0.1 | 0.1 | 0.1 |
| *Phaseolus perplexus* | 1 | 0.1 | 0.1 | 0.1 | 0.1 | 0.1 | 0.1 |
| *Phaseolus pluriflorus* | 1 | 0.5 | 0.1 | 0.1 | 0.1 | 0.5 | 0.1 |
| *Phaseolus ritensis* | 1 | 1 | 0.1 | 0.1 | 0.1 | 0.1 | 0.1 |
| *Phaseolus scabrellus* | 1 | 1 | 0.1 | 0.1 | 0.1 | 0.1 | 0.1 |
| *Phaseolus vulgaris* | 0.1 | 1 | 0.5 | 1 | 0.1 | 1 | 0.5 |
| *Physalis acutifolia* | 1 | 1 | 1 | 1 | 1 | 0.1 | 0.1 |
| *Physalis angulata* | 1 | 1 | 1 | 1 | 0.1 | 0.1 | 0.1 |
| *Physalis angustiphysa* | 1 | 0.1 | 0.1 | 0.5 | 0.1 | 0.1 | 0.1 |
| *Physalis campechiana* | 1 | 1 | 0.1 | 0.1 | 0.1 | 0.1 | 0.1 |
| *Physalis caudella* | 1 | 1 | 0.1 | 0.1 | 0.1 | 0.1 | 0.1 |
| *Physalis chenopodiifolia* | 1 | 1 | 0.1 | 0.1 | 0.1 | 0.5 | 0.1 |
| *Physalis cinerascens* | 1 | 1 | 0.1 | 0.1 | 0.1 | 1 | 0.1 |
| *Physalis cordata* | 1 | 1 | 0.1 | 1 | 0.1 | 0.1 | 0.1 |
| *Physalis coztomatl* | 1 | 0.1 | 0.1 | 0.1 | 0.1 | 0.1 | 0.1 |
| *Physalis crassifolia* | 1 | 1 | 0.1 | 0.1 | 0.1 | 0.1 | 0.1 |
| *Physalis glutinosa* | 1 | 1 | 0.1 | 0.1 | 0.1 | 0.1 | 0.1 |
| *Physalis gracilis* | 1 | 1 | 0.1 | 0.5 | 0.1 | 0.5 | 0.1 |
| *Physalis hederifolia* | 1 | 1 | 0.1 | 0.1 | 0.1 | 0.1 | 0.1 |
| *Physalis ignota* | 1 | 1 | 0.1 | 0.1 | 0.1 | 0.1 | 0.1 |
| *Physalis lagascae* | 1 | 1 | 0.1 | 1 | 0.1 | 0.1 | 1 |
| *Physalis leptophylla* | 1 | 1 | 0.1 | 0.1 | 0.1 | 0.1 | 0.1 |
| *Physalis melanocystis* | 1 | 1 | 0.1 | 0.1 | 0.1 | 0.1 | 0.1 |
| *Physalis minuta* | 1 | 0.1 | 0.1 | 0.1 | 0.1 | 0.1 | 0.1 |
| *Physalis nicandroides* | 1 | 1 | 0.1 | 1 | 0.1 | 1 | 0.1 |
| *Physalis orizabae* | 1 | 0.1 | 0.1 | 0.1 | 0.1 | 0.1 | 0.1 |
| *Physalis patula* | 1 | 1 | 0.1 | 1 | 0.1 | 0.1 | 1 |
| *Physalis philadelphica* | 1 | 1 | 1 | 1 | 0.1 | 0.1 | 1 |
| *Physalis pruinosa* | 1 | 1 | 0.1 | 1 | 0.1 | 0.1 | 0.1 |
| *Physalis pubescens* | 1 | 1 | 0.1 | 1 | 0.1 | 0.1 | 0.1 |

| | | | | | | | |
|---|---|---|---|---|---|---|---|
| *Physalis solanacea* | 1 | 1 | 0.1 | 1 | 0.1 | 0.1 | 0.1 |
| *Physalis sordida* | 1 | 1 | 0.1 | 0.1 | 0.1 | 0.1 | 0.1 |
| *Physalis sulphurea* | 1 | 1 | 0.1 | 0.1 | 0.1 | 0.1 | 0.1 |
| *Physalis virginiana* | 1 | 1 | 0.1 | 1 | 0.1 | 0.1 | 0.1 |
| *Physalis volubilis* | 1 | 0.1 | 0.1 | 0.1 | 0.1 | 0.1 | 0.1 |
| *Physalis waterfallii* | 1 | 0.1 | 0.1 | 0.1 | 0.1 | 0.1 | 0.1 |
| *Solanum agrimonifolium* | 1 | 1 | 1 | 0.1 | 0.1 | 0.1 | 0.1 |
| *Solanum bulbocastanum* | 1 | 1 | 1 | 0.1 | 0.1 | 0.1 | 0.1 |
| *Solanum cardiophyllum* | 1 | 1 | 1 | 0.1 | 0.1 | 0.1 | 0.1 |
| *Solanum clarum* | 1 | 1 | 0.1 | 0.1 | 0.1 | 0.1 | 0.1 |
| *Solanum demissum* | 1 | 1 | 1 | 0.1 | 0.1 | 0.1 | 0.1 |
| *Solanum ehrenbergii* | 1 | 1 | 1 | 0.1 | 0.1 | 0.1 | 0.1 |
| *Solanum hjertingii* | 1 | 1 | 1 | 0.1 | 0.1 | 0.1 | 0.1 |
| *Solanum hougasii* | 1 | 1 | 1 | 0.1 | 0.1 | 0.1 | 0.1 |
| *Solanum iopetalum* | 1 | 1 | 1 | 0.1 | 0.1 | 0.1 | 0.1 |
| *Solanum morelliforme* | 1 | 1 | 0.1 | 0.1 | 0.1 | 0.1 | 0.1 |
| *Solanum oxycarpum* | 1 | 1 | 1 | 0.1 | 0.1 | 0.1 | 0.1 |
| *Solanum pinnatisectum* | 1 | 1 | 1 | 0.1 | 0.1 | 0.1 | 0.1 |
| *Solanum polyadenium* | 1 | 1 | 1 | 0.1 | 0.1 | 0.1 | 0.1 |
| *Solanum schenckii* | 1 | 1 | 1 | 0.1 | 0.1 | 0.1 | 0.1 |
| *Solanum stenophyllidium* | 1 | 1 | 1 | 0.1 | 0.1 | 0.1 | 0.1 |
| *Solanum stoloniferum* | 1 | 1 | 1 | 0.1 | 0.1 | 0.1 | 0.1 |
| *Solanum trifidum* | 1 | 1 | 1 | 0.1 | 0.1 | 0.1 | 0.1 |
| *Solanum verrucosum* | 1 | 1 | 1 | 0.1 | 0.1 | 0.1 | 0.1 |
| *Tripsacum lanceolatum* | 1 | 1 | 0.1 | 0.1 | 0.1 | 0.1 | 0.1 |
| *Tripsacum latifolium* | 1 | 1 | 0.1 | 0.1 | 0.1 | 1 | 0.1 |
| *Tripsacum pilosum* | 1 | 1 | 0.1 | 0.1 | 0.1 | 0.1 | 0.1 |
| *Vanilla odorata* | 1 | 1 | 1 | 0.5 | 0.1 | 0.1 | 0.1 |
| *Vanilla pompona* | 1 | 1 | 1 | 1 | 0.1 | 0.1 | 0.1 |
| *Zea diploperennis* | 0.1 | 1 | 0.1 | 1 | 0.1 | 1 | 0.1 |
| *Zea luxurians* | 0.1 | 1 | 0.1 | 1 | 1 | 1 | 0.1 |
| *Zea mays* subsp. *huehuetenangensis* | 0.1 | 1 | 0.1 | 0.1 | 0.1 | 0.1 | 0.1 |
| *Zea mays* subsp. *mexicana* Chalco subpopulation | 0.1 | 1 | 1 | 1 | 0.1 | 0.1 | 0.1 |
| *Zea mays* subsp. *mexicana* Durango subpopulation | 0.1 | 1 | 1 | 1 | 0.1 | 0.1 | 0.1 |
| *Zea mays* subsp. *mexicana* Mesa-Central subpopulation | 0.1 | 1 | 1 | 1 | 1 | 0.1 | 0.1 |
| *Zea mays* subsp. *mexicana* Nobogame subpopulation | 0.1 | 1 | 1 | 1 | 0.1 | 0.1 | 0.1 |
| *Zea mays* subsp. *parviglumis* | 0.1 | 1 | 1 | 1 | 1 | 0.1 | 0.5 |
| *Zea perennis* | 0.1 | 1 | 0.5 | 1 | 0.1 | 0.1 | 0.1 |

**Supplementary Table 8. MaxEnt performance and significance of potential species distribution models (SDM). (Occurrence data was divided into 70% for training and 30% for testing the model. MaxEnt performance was tested with 30% of the withheld data. Testing omission rate is given for SDM selected by experts, which were mainly model threshold with the ten percentile training presence.) (*indicates omission rates under a minimum training presence threshold.)**

| Taxa | Training records | Testing records | AUC value | Testing omission rate* |
|---|---|---|---|---|
| *Capsicum annuum* | 205 | 88 | 0.81 | 0.13 |
| *Capsicum frutescens* | 282 | 121 | 0.77 | 0.18 |
| *Capsicum lanceolatum* | 37 | 17 | 0.9 | 0.06 |
| *Capsicum rhomboideum* | 157 | 68 | 0.85 | 0.16 |
| *Cucurbita argyrosperma* | 249 | 108 | 0.84 | 0.14 |
| *Cucurbita cordata* | 58 | 26 | 0.74 | 0.12 |
| *Cucurbita digitata* | 77 | 34 | 0.87 | 0.09 |
| *Cucurbita foetidissima* | 585 | 251 | 0.82 | 0.13 |
| *Cucurbita lundelliana* | 32 | 14 | 0.73 | 0.21 |
| *Cucurbita okeechobeensis* subsp. *martinezii* | 72 | 31 | 0.9 | 0.19 |
| *Cucurbita palmata* | 189 | 82 | 0.91 | 0.23 |
| *Cucurbita pedatifolia* | 44 | 20 | 0.87 | 0.15 |
| *Cucurbita pepo* subsp. *fraterna* | 20 | 9 | 0.75 | 0.33 |
| *Cucurbita radicans* | 54 | 24 | 0.89 | 0.17 |
| *Gossypium aridum* | 103 | 45 | 0.92 | 0* |
| *Gossypium davidsonii* | 35 | 15 | 0.93 | 0.14 |
| *Gossypium gossypioides* | 23 | 10 | 0.98 | 0.1 |
| *Gossypium harknessii* | 39 | 18 | 0.95 | 0.07 |
| *Gossypium hirsutum* | 157 | 68 | 0.85 | 0.05* |
| *Gossypium thurberi* | 62 | 28 | 0.93 | 0.11 |
| *Persea albida* | 16 | 8 | 0.74 | 0.63 |
| *Persea americana* | 49 | 21 | 0.8 | 0 |
| *Persea caerulea* | 214 | 92 | 0.9 | 0.1 |
| *Persea donnell-smithii* | 42 | 18 | 0.81 | 0.28 |
| *Persea hintonii* | 56 | 25 | 0.96 | 0 |
| *Persea liebmannii* | 119 | 52 | 0.91 | 0.12 |
| *Persea longipes* | 23 | 10 | 0.91 | 0.1 |
| *Persea pallescens* | 14 | 7 | 0.81 | 0.29 |
| *Persea podadenia* | 17 | 8 | 0.78 | 0.13 |

| | | | | |
|---|---|---|---|---|
| *Persea purpusii* | 25 | 11 | 0.98 | 0.09 |
| *Persea schiedeana* | 118 | 52 | 0.93 | 0.06 |
| *Persea vesticula* | 23 | 11 | 0.94 | 0 |
| *Physalis acutifolia* | 113 | 49 | 0.88 | 0.1 |
| *Physalis angulata* | 277 | 120 | 0.75 | 0.12 |
| *Physalis angustiphysa* | 31 | 14 | 0.84 | 0* |
| *Physalis campechiana* | 18 | 8 | 0.74 | 0.25 |
| *Physalis caudella* | 32 | 14 | 0.78 | 0.31 |
| *Physalis chenopodiifolia* | 97 | 42 | 0.89 | 0.14 |
| *Physalis cinerascens* | 118 | 52 | 0.85 | 0.25 |
| *Physalis cordata* | 89 | 39 | 0.84 | 0.14 |
| *Physalis coztomatl* | 74 | 33 | 0.9 | 0.06 |
| *Physalis crassifolia* | 503 | 214 | 0.96 | 0.1 |
| *Physalis glutinosa* | 28 | 13 | 0.84 | 0.15 |
| *Physalis gracilis* | 119 | 52 | 0.77 | 0.15 |
| *Physalis hederifolia* | 171 | 74 | 0.82 | 0* |
| *Physalis ignota* | 65 | 29 | 0.96 | |
| *Physalis lagascae* | 140 | 60 | 0.86 | 0.17 |
| *Physalis leptophylla* | 59 | 26 | 0.87 | 0.12 |
| *Physalis melanocystis* | 37 | 17 | 0.83 | 0.06 |
| *Physalis minuta* | 23 | 11 | 0.83 | 0* |
| *Physalis nicandroides* | 159 | 69 | 0.86 | 0.13 |
| *Physalis orizabae* | 146 | 63 | 0.82 | 0.17 |
| *Physalis patula* | 100 | 44 | 0.87 | 0* |
| *Physalis philadelphica* | 359 | 155 | 0.89 | 0.1 |
| *Physalis pruinosa* | 116 | 50 | 0.88 | 0.2 |
| *Physalis pubescens* | 421 | 181 | 0.78 | 0.09 |
| *Physalis solanacea* | 58 | 25 | 0.75 | 0.2* |
| *Physalis sordida* | 22 | 10 | 0.84 | 0* |
| *Physalis sulphurea* | 37 | 16 | 0.91 | 0.19 |
| *Physalis virginiana* | 43 | 19 | 0.83 | 0.11 |
| *Physalis volubilis* | 49 | 22 | 0.92 | 0.14 |
| *Physalis waterfallii* | 20 | 10 | 0.92 | 0.1 |
| *Solanum agrimonifolium* | 34 | 15 | 0.97 | 0 |
| *Solanum bulbocastanum* | 214 | 93 | 0.91 | 0.13 |
| *Solanum cardiophyllum* | 111 | 48 | 0.9 | 0.17 |
| *Solanum clarum* | 23 | 11 | 0.95 | 0.27 |
| *Solanum demissum* | 274 | 118 | 0.97 | 0.14 |
| *Solanum ehrenbergii* | 84 | 36 | 0.85 | 0.25 |
| *Solanum hjertingii* | 35 | 16 | 0.94 | 0.13 |
| *Solanum hougasii* | 49 | 22 | 0.92 | 0.18 |

| | | | | |
|---|---|---|---|---|
| *Solanum iopetalum* | 184 | 80 | 0.94 | 0.06 |
| *Solanum morelliforme* | 68 | 30 | 0.88 | 0.13 |
| *Solanum oxycarpum* | 49 | 10 | 0.94 | 0.11 |
| *Solanum pinnatisectum* | 49 | 21 | 0.86 | 0.14 |
| *Solanum polyadenium* | 56 | 24 | 0.84 | 0.17 |
| *Solanum schenckii* | 41 | 18 | 0.94 | 0.11 |
| *Solanum stenophyllidium* | 70 | 31 | 0.87 | 0.16 |
| *Solanum stoloniferum* | 691 | 297 | 0.9 | 0.15 |
| *Solanum trifidum* | 63 | 28 | 0.91 | 0.21 |
| *Solanum verrucosum* | 174 | 75 | 0.93 | 0.17 |
| *Tripsacum lanceolatum* | 142 | 62 | 0.81 | 0.18 |
| *Tripsacum latifolium* | 25 | 11 | 0.7 | 0.27 |
| *Tripsacum pilosum* | 27 | 12 | 0.87 | 0.25 |
| *Vanilla odorata* | 28 | 12 | 0.69 | 0.13 |
| *Vanilla pompona* | 17 | 10 | 0.79 | 0.11 |

**Supplementary Table 9. Direct download links for CWR potential species distribution models (SDM).**

| Title of potential SDM | Direct download link (available upon aceptance) |
|---|---|
| Capsicum annuum var. glabriusculum Distribución potencial de chiles | |
| Capsicum frutescens Distribución potencial de chiles | |
| Capsicum lanceolatum Distribución potencial de chiles | |
| Capsicum rhomboideum Distribución potencial de chiles | |
| Cucurbita argyrosperma subsp. sororia Distribución potencial de calabazas | |
| Cucurbita cordata Distribución potencial de calabazas | |
| Cucurbita digitata Distribución potencial de calabazas | |
| Cucurbita foetidissima Distribución potencial de calabazas | |
| Cucurbita lundelliana Distribución potencial de calabazas | |
| Cucurbita okeechobeensis subsp. martinezii Distribución potencial de calabazas | |
| Cucurbita palmata Distribución potencial de calabazas | |
| Cucurbita pedatifolia Distribución potencial de calabazas | |
| Cucurbita pepo subsp. fraterna Distribución potencial de calabazas | |
| Cucurbita radicans Distribución potencial de calabazas | |
| Gossypium aridum Distribución potencial de algodones | |
| Gossypium davidsonii Distribución potencial de algodones | |
| Gossypium gossypioides Distribución potencial de algodones | |
| Gossypium harknesii Distribución potencial de algodones | |
| Gossypium hirsutum Distribución potencial de algodones | |
| Gossypium thurberi Distribución potencial de algodones | |
| Persea albida Distribución potencial de aguacates | |
| Persea americana Distribución potencial de aguacates | |
| Persea caerulea Distribución potencial de aguacates | |
| Persea donnell-smithii Distribución potencial de aguacates | |

| |
|---|
| Persea hintonii Distribución potencial de aguacates |
| Persea liebmannii Distribución potencial de aguacates |
| Persea longipes Distribución potencial de aguacates |
| Persea pallescens Distribución potencial de aguacates |
| Persea podadenia Distribución potencial de aguacates |
| Persea purpusii Distribución potencial de aguacates |
| Persea schiedeana Distribución potencial de aguacates |
| Persea vesticula Distribución potencial de aguacates |
| Physalis acutifolia Distribución potencial de tomates verdes |
| Physalis angulata Distribución potencial de tomates verdes |
| Physalis angustiphysa Distribución potencial de tomates verdes |
| Physalis campechiana Distribución potencial de tomates verdes |
| Physalis caudella Distribución potencial de tomates verdes |
| Physalis cinerascens Distribución potencial de tomates verdes |
| Physalis cordata Distribución potencial de tomates verdes |
| Physalis coztomatl Distribución potencial de tomates verdes |
| Physalis crassifolia Distribución potencial de tomates verdes |
| Physalis chenopodiifolia Distribución potencial de tomates verdes |
| Physalis glutinosa Distribución potencial de tomates verdes |
| Physalis gracilis Distribución potencial de tomates verdes |
| Physalis hederifolia Distribución potencial de tomates verdes |
| Physalis ignota Distribución potencial de tomates verdes |
| Physalis lagascae Distribución potencial de tomates verdes |
| Physalis leptophylla Distribución potencial de tomates verdes |
| Physalis melanocystis Distribución potencial de tomates verdes |
| Physalis minuta Distribución potencial de tomates verdes |
| Physalis nicandroides Distribución potencial de tomates verdes |
| Physalis orizabae Distribución potencial de tomates verdes |
| Physalis patula Distribución potencial de tomates verdes |

| |
|---|
| Physalis philadelphica Distribución potencial de tomates verdes |
| Physalis pruinosa Distribución potencial de tomates verdes |
| Physalis pubescens Distribución potencial de tomates verdes |
| Physalis solanaceus Distribución potencial de tomates verdes |
| Physalis sordida Distribución potencial de tomates verdes |
| Physalis sulphurea Distribución potencial de tomates verdes |
| Physalis virginiana Distribución potencial de tomates verdes |
| Physalis volubilis Distribución potencial de tomates verdes |
| Physalis waterfallii Distribución potencial de tomates verdes |
| Solanum agrimonifolium Distribución potencial de papas |
| Solanum bulbocastanum Distribución potencial de papas |
| Solanum cardiophyllum Distribución potencial de papas |
| Solanum clarum Distribución potencial de papas |
| Solanum demissum Distribución potencial de papas |
| Solanum ehrenbergii Distribución potencial de papas |
| Solanum hjertingii Distribución potencial de papas |
| Solanum hougasii Distribución potencial de papas |
| Solanum iopetalum Distribución potencial de papas |
| Solanum morelliforme Distribución potencial de papas |
| Solanum oxycarpum Distribución potencial de papas |
| Solanum pinnatisectum Distribución potencial de papas |
| Solanum polyadenium Distribución potencial de papas |
| Solanum schenckii Distribución potencial de papas |
| Solanum stenophyllidium Distribución potencial de papas |
| Solanum stoloniferum Distribución potencial de papas |
| Solanum trifidum Distribución potencial de papas |
| Solanum verrucosum Distribución potencial de papas |
| Tripsacum lanceolatum Distribución potencial de maíces |
| Tripsacum latifolium Distribución potencial de maíces |

| |
|---|
| Tripsacum pilosum Distribución potencial de maíces |
| Vanilla odorata Distribución potencial de vainillas |
| Vanilla pompona Distribución potencial de vainillas |

**Supplementary Table 10. CWR in protected areas. Information is based on occurrences (occs.) of 224 taxa and 116 potential species distribution models (SDM).**

| No. | ID_Protected area | NAME | Official extension ($km^2$) | # occs | # taxa based on occs | # taxa based on SDM |
|---|---|---|---|---|---|---|
| 1 | 2.1.01.104 | Alto Golfo de California y Delta del Río Colorado | 9347.6 | 6 | 2 | 11 |
| 2 | 9.3.03.111 | Arrecife Alacranes | 3337.7 | NA | NA | NA |
| 3 | 9.3.07.128 | Arrecife de Puerto Morelos | 90.7 | NA | NA | NA |
| 4 | 9.3.05.123 | Arrecifes de Cozumel | 119.9 | NA | NA | NA |
| 5 | 9.1.04.127 | Arrecifes de Sian Ka'an | 349.3 | NA | NA | NA |
| 6 | 9.3.08.145 | Arrecifes de Xcalak | 179.5 | NA | NA | NA |
| 7 | 1.3.04.122 | Bahía de Loreto | 2065.8 | 27 | 4 | 5 |
| 8 | 9.7.04.159 | Bala'an K'aax | 1283.9 | 0 | 0 | 17 |
| 9 | 1.7.03.176 | Balandra | 25.1 | 1 | 1 | 6 |
| 10 | 9.1.03.124 | Banco Chinchorro | 1443.6 | NA | NA | NA |
| 11 | 6.1.05.144 | Barranca de Metztitlán | 960.4 | 10 | 7 | 52 |
| 12 | 5.3.03.033 | Barranca del Cupatitzio | 4.6 | 1 | 1 | 46 |
| 13 | 2.7.04.182 | Bavispe | 2009 | 20 | 5 | 28 |
| 14 | 8.3.02.023 | Benito Juárez | 25.9 | 4 | 4 | 45 |
| 15 | 8.4.01.099 | Bonampak | 43.6 | 0 | 0 | 21 |
| 16 | 8.7.05.166 | Boquerón de Tonalá | 39.1 | 1 | 1 | 38 |
| 17 | 6.3.20.040 | Bosencheve | 146 | 22 | 6 | 52 |
| 18 | 5.6.02.149 | C.A.D.N.R. 001 Pabellón | 977 | 12 | 5 | 48 |
| 19 | 4.6.01.150 | C.A.D.N.R. 004 Don Martín | 15193.9 | 31 | 5 | 29 |
| 20 | 4.6.02.151 | C.A.D.N.R. 026 Bajo Río San Juan | 1971.6 | 29 | 5 | 40 |
| 21 | 5.6.03.152 | C.A.D.N.R. 043 Estado de Nayarit | 23290.3 | 192 | 46 | 83 |
| 22 | 4.7.03.113 | Cañón de Santa Elena | 2772.1 | 1 | 1 | 26 |
| 23 | 7.3.03.025 | Cañón del Río Blanco | 488 | 72 | 23 | 62 |
| 24 | 8.3.04.060 | Cañón del Sumidero | 217.9 | 25 | 14 | 47 |
| 25 | 7.7.03.165 | Cañon del Usumacinta | 461.3 | 0 | 0 | 24 |
| 26 | 1.3.03.117 | Cabo Pulmo | 71.1 | 0 | 0 | 5 |
| 27 | 1.7.01.050 | Cabo San Lucas | 40 | 2 | 1 | 0 |
| 28 | 9.1.02.095 | Calakmul | 7231.9 | 21 | 13 | 20 |
| 29 | 3.7.02.024 | Campo Verde | 1080.7 | 0 | 0 | 24 |
| 30 | 9.1.09.178 | Caribe Mexicano | 57540.6 | NA | NA | NA |
| 31 | 8.7.01.057 | Cascada de Agua Azul | 25.8 | 0 | 0 | 19 |
| 32 | 3.3.02.062 | Cascada de Bassaseachic | 58 | 18 | 5 | 30 |
| 33 | 5.3.01.005 | Cerro de Garnica | 19.4 | 7 | 4 | 47 |
| 34 | 6.3.16.028 | Cerro de La Estrella | 11.8 | 1 | 1 | 40 |
| 35 | 4.4.01.097 | Cerro de la Silla | 60.4 | 7 | 2 | 26 |
| 36 | 6.3.11.019 | Cerro de Las Campanas | 0.6 | 0 | 0 | 30 |

| | | | | | | |
|---|---|---|---|---|---|---|
| 37 | 3.7.05.177 | Cerro Mohinora | 91.3 | 4 | 3 | 28 |
| 38 | 5.1.02.106 | Chamela-Cuixmala | 131.4 | 30 | 13 | 29 |
| 39 | 8.7.02.100 | Chan-Kin | 121.8 | 0 | 0 | 16 |
| 40 | 6.7.03.153 | Ciénegas del Lerma | 30.2 | 3 | 3 | 48 |
| 41 | 7.3.02.017 | Cofre de Perote o Nauhcampatépetl | 115.3 | 17 | 4 | 39 |
| 42 | 1.1.01.049 | Complejo Lagunar Ojo de Liebre | 793.3 | 0 | 0 | 6 |
| 43 | 1.3.02.047 | Constitución de 1857 | 50.1 | 0 | 0 | 3 |
| 44 | 6.7.02.093 | Corredor Biológico Chichinautzin | 373 | 82 | 17 | 67 |
| 45 | 9.3.04.120 | Costa Occ. de I. Mujeres, Pta. Cancún y Pta. Nizuc | 86.7 | 1 | 1 | 0 |
| 46 | 4.7.05.115 | Cuatrociénegas | 843.5 | 0 | 0 | 11 |
| 47 | 3.3.01.037 | Cumbres de Majalca | 47 | 22 | 8 | 24 |
| 48 | 4.3.05.139 | Cumbres de Monterrey | 1774 | 66 | 14 | 40 |
| 49 | 6.3.06.011 | Cumbres del Ajusco | 9.2 | 21 | 7 | 35 |
| 50 | 6.3.01.001 | Desierto de los Leones | 15.3 | 35 | 4 | 42 |
| 51 | 6.3.21.043 | Desierto del Carmen o de Nixcongo | 5.3 | 0 | 0 | 49 |
| 52 | 9.3.02.091 | Dzibilchantún | 5.4 | 7 | 2 | 13 |
| 53 | 6.3.25.069 | El Chico | 27.4 | 46 | 8 | 33 |
| 54 | 6.3.26.070 | El Cimatario | 24.5 | 3 | 2 | 44 |
| 55 | 6.3.17.030 | El Histórico Coyoacán | 0.4 | 4 | 2 | 0 |
| 56 | 5.7.03.068 | El Jabalí | 51.8 | 40 | 14 | 64 |
| 57 | 2.1.02.105 | El Pinacate y Gran Desierto de Altar | 7145.6 | 57 | 6 | 9 |
| 58 | 4.3.01.008 | El Potosí | 20 | 0 | 0 | 32 |
| 59 | 4.3.03.029 | El Sabinal | 0.1 | 0 | 0 | 0 |
| 60 | 6.3.10.016 | El Tepeyac | 15 | 3 | 2 | 40 |
| 61 | 6.3.09.015 | El Tepozteco | 232.6 | 57 | 12 | 66 |
| 62 | 8.1.03.096 | El Triunfo | 1191.8 | 30 | 9 | 42 |
| 63 | 6.3.23.059 | El Veladero | 36.2 | 0 | 0 | 26 |
| 64 | 1.1.02.094 | El Vizcaíno | 25467.9 | 45 | 4 | 13 |
| 65 | 6.3.07.012 | Fuentes Brotantes de Tlalpan | 1.3 | 0 | 0 | 0 |
| 66 | 6.3.22.048 | General Juan Álvarez | 5.3 | 0 | 0 | 30 |
| 67 | 4.3.02.010 | Gogorrón | 380.1 | 4 | 2 | 40 |
| 68 | 6.3.03.004 | Grutas de Cacahuamilpa | 16 | 1 | 1 | 36 |
| 69 | 8.3.06.129 | Huatulco | 118.9 | 19 | 3 | 25 |
| 70 | 5.3.04.034 | Insurgente José María Morelos | 71.9 | 15 | 5 | 62 |
| 71 | 6.3.05.009 | Insurgente Miguel Hidalgo y Costilla | 18.9 | 3 | 3 | 43 |
| 72 | 9.3.06.126 | Isla Contoy | 51.3 | ND | ND | ND |
| 73 | 1.1.05.157 | Isla Guadalupe | 4769.7 | ND | ND | ND |
| 74 | 5.3.07.061 | Isla Isabel | 1.9 | ND | ND | ND |
| 75 | 2.1.03.147 | Isla San Pedro Mártir | 301.7 | ND | ND | ND |
| 76 | 2.7.01.052 | Islas del Golfo de California | 3745.5 | 67 | 6 | ND |
| 77 | 1.1.07.179 | Islas del Pacífico de la Península de Baja California | 11612.2 | 36 | 4 | ND |

| | | | | | | |
|---|---|---|---|---|---|---|
| 78 | 5.8.08.148 | Islas La Pajarera, Cocinas, Mamut, Colorada, San Pedro, San Agustín, San Andrés y Negrita y los Islotes Los Anegados, Novillas, Mosca y Submarino | 19.8 | 5 | 4 | ND |
| 79 | 5.1.03.142 | Islas Marías | 6412.8 | 1 | 1 | ND |
| 80 | 5.3.08.158 | Islas Marietas | 13.8 | ND | ND | ND |
| 81 | 6.3.02.002 | Iztaccíhuatl-Popocatépetl | 398.2 | 94 | 7 | 47 |
| 82 | 3.1.02.173 | Janos | 5264.8 | 13 | 5 | 24 |
| 83 | 8.1.05.118 | La Encrucijada | 1448.7 | 0 | 0 | 12 |
| 84 | 3.1.01.054 | La Michilía | 350 | 7 | 3 | 35 |
| 85 | 6.3.18.031 | La Montaña Malinche o Matlalcuéyatl | 461.1 | 57 | 6 | 52 |
| 86 | 9.7.06.175 | La porción norte y la franja costera oriental, terrestres y marinas de la Isla de Cozumel | 378.3 | NA | NA | NA |
| 87 | 5.7.02.056 | La Primavera | 305 | 37 | 19 | 58 |
| 88 | 8.1.06.119 | La Sepultura | 1673.1 | 32 | 12 | 57 |
| 89 | 8.1.04.101 | Lacan-Tun | 618.7 | 0 | 0 | 24 |
| 90 | 5.3.05.041 | Lago de Camácuaro | 0.1 | NA | NA | NA |
| 91 | 7.7.01.112 | Laguna de Términos | 7061.5 | 11 | 4 | 16 |
| 92 | 7.7.02.155 | Laguna Madre y Delta del Río Bravo | 5728.1 | 3 | 3 | 23 |
| 93 | 8.3.01.020 | Lagunas de Chacahua | 149 | 1 | 1 | 26 |
| 94 | 8.3.03.046 | Lagunas de Montebello | 64.3 | 18 | 8 | 30 |
| 95 | 6.3.08.013 | Lagunas de Zempoala | 47.9 | 44 | 8 | 45 |
| 96 | 5.6.01.092 | Las Huertas | 1.7 | 2 | 1 | 28 |
| 97 | 6.3.15.027 | Lomas de Padierna | 11.6 | 14 | 7 | 46 |
| 98 | 6.3.04.007 | Los Mármoles | 231.5 | 43 | 9 | 45 |
| 99 | 4.3.04.038 | Los Novillos | 0.4 | 0 | 0 | 3 |
| 100 | 9.1.06.136 | Los Petenes | 2828.6 | 2 | 2 | 18 |
| 101 | 6.3.14.026 | Los Remedios | 4 | 0 | 0 | 44 |
| 102 | 7.1.02.133 | Los Tuxtlas | 1551.2 | 39 | 12 | 14 |
| 103 | 3.7.04.167 | Médanos de Samalayuca | 631.8 | 0 | 0 | 11 |
| 104 | 4.7.04.114 | Maderas del Carmen | 2083.8 | 18 | 4 | 26 |
| 105 | 9.7.05.164 | Manglares de Nichupté | 42.6 | 0 | 0 | 12 |
| 106 | 4.1.01.055 | Mapimí | 3423.9 | 10 | 3 | 15 |
| 107 | 6.1.04.138 | Mariposa Monarca | 562.6 | 63 | 13 | 59 |
| 108 | 5.1.05.174 | Marismas Nacionales Nayarit | 1338.5 | 3 | 1 | 31 |
| 109 | 2.7.03.140 | Meseta de Cacaxtla | 508.6 | 2 | 1 | 30 |
| 110 | 8.7.04.132 | Metzabok | 33.7 | 0 | 0 | 22 |
| 111 | 6.3.12.021 | Molino de Flores Netzahualcóyotl | 0.5 | 0 | 0 | 0 |
| 112 | 8.1.01.051 | Montes Azules | 3312 | 21 | 9 | 32 |
| 113 | 8.7.03.131 | Nahá | 38.5 | 3 | 2 | 25 |
| 114 | 6.7.01.003 | Nevado de Toluca | 535.9 | 112 | 6 | 48 |
| 115 | 4.7.06.168 | Ocampo | 3442.4 | 2 | 1 | 14 |

| | | | | | | |
|---|---|---|---|---|---|---|
| 116 | 9.7.03.146 | Otoch Ma'ax Yetel Kooh | 53.7 | 0 | 0 | 12 |
| 117 | 1.1.08.180 | Pacífico Mexicano Profundo | 436141.2 | NA | NA | NA |
| 118 | 8.3.05.066 | Palenque | 17.7 | 0 | 0 | 16 |
| 119 | 7.1.01.098 | Pantanos de Centla | 3027.1 | 1 | 1 | 12 |
| 120 | 3.7.03.035 | Papigochic | 2227.6 | 20 | 4 | 30 |
| 121 | 7.3.01.014 | Pico de Orizaba | 197.5 | 18 | 4 | 38 |
| 122 | 5.7.01.039 | Pico de Tancítaro | 234.1 | 24 | 7 | 52 |
| 123 | 9.8.01.085 | Playa adyacente a la localidad denominada RÍo Lagartos | 6.1 | 0 | 0 | 0 |
| 124 | 2.8.01.078 | Playa Ceuta | 1.4 | 0 | 0 | 0 |
| 125 | 5.8.05.082 | Playa Cuitzmala | 0.2 | 0 | 0 | 2 |
| 126 | 8.8.03.087 | Playa de Escobilla | 1.5 | 0 | 0 | 0 |
| 127 | 8.8.04.089 | Playa de la BahÍa de Chacahua | 0.9 | 0 | 0 | 6 |
| 128 | 9.8.02.086 | Playa de la Isla Contoy | 0.1 | 0 | 0 | ND |
| 129 | 5.8.02.075 | Playa de Maruata y Colola | 2.2 | 0 | 0 | 0 |
| 130 | 5.8.06.084 | Playa de Mismaloya | 6.3 | 0 | 0 | 6 |
| 131 | 8.8.02.083 | Playa de Puerto Arista | 2.1 | 0 | 0 | 2 |
| 132 | 7.8.01.076 | Playa de Rancho Nuevo | 0.9 | 0 | 0 | 0 |
| 133 | 8.8.01.080 | Playa de Tierra Colorada | 1.4 | 0 | 0 | 0 |
| 134 | 5.8.07.088 | Playa El Tecuán | 0.4 | 0 | 0 | 0 |
| 135 | 2.8.02.079 | Playa El Verde Camacho | 1 | 0 | 0 | 0 |
| 136 | 5.8.01.074 | Playa Mexiquillo | 0.7 | 0 | 0 | 0 |
| 137 | 5.8.03.077 | Playa Piedra de Tlacoyunque | 1 | 0 | 0 | 0 |
| 138 | 5.8.04.081 | Playa Teopa | 0.3 | 0 | 0 | 0 |
| 139 | 1.3.07.107 | Revillagigedo | 148087.8 | 12 | 3 | 4 |
| 140 | 9.1.07.143 | Ría Celestún | 814.8 | 7 | 1 | 15 |
| 141 | 9.1.05.134 | Ría Lagartos | 603.5 | 11 | 4 | 15 |
| 142 | 4.4.02.172 | Río Bravo del Norte | 21.8 | 0 | 0 | 6 |
| 143 | 5.3.06.045 | Rayón | 0.3 | 0 | 0 | 0 |
| 144 | 6.3.19.036 | Sacromonte | 0.4 | 6 | 2 | 0 |
| 145 | 8.1.02.072 | Selva El Ocote | 1012.9 | 10 | 4 | 42 |
| 146 | 9.1.01.073 | Sian Ka'an | 5281.5 | 5 | 3 | 15 |
| 147 | 3.3.03.141 | Sierra de Árganos | 11.2 | 0 | 0 | 26 |
| 148 | 2.7.02.121 | Sierra de Álamos-Río Cuchujaqui | 928.9 | 63 | 10 | 39 |
| 149 | 4.7.01.063 | Sierra de Álvarez | 169 | 23 | 6 | 40 |
| 150 | 6.1.03.137 | Sierra de Huautla | 590.3 | 6 | 6 | 60 |
| 151 | 5.1.01.090 | Sierra de Manantlán | 1395.8 | 255 | 42 | 72 |
| 152 | 5.7.04.071 | Sierra de Quila | 151.9 | 6 | 4 | 65 |
| 153 | 1.3.01.044 | Sierra de San Pedro MÃ¡rtir | 729.1 | 0 | 0 | 7 |
| 154 | 7.1.03.181 | Sierra de Tamaulipas | 3088.9 | 17 | 6 | 36 |
| 155 | 4.1.02.108 | Sierra del Abra Tanchipa | 214.6 | 0 | 0 | 30 |
| 156 | 6.1.01.125 | Sierra Gorda | 3835.7 | 215 | 32 | 58 |
| 157 | 6.1.06.160 | Sierra Gorda de Guanajuato | 2368.8 | 54 | 22 | 61 |
| 158 | 1.1.04.109 | Sierra La Laguna | 1124.4 | 25 | 7 | 11 |
| 159 | 4.7.02.067 | Sierra La Mojonera | 92 | 0 | 0 | 10 |

| | | | | | | |
|---|---|---|---|---|---|---|
| 160 | 7.7.04.169 | Sistema Arrecifal Lobos-Tuxpan | 305.7 | NA | NA | NA |
| 161 | 7.3.04.103 | Sistema Arrecifal Veracruzano | 655.2 | NA | NA | NA |
| 162 | 6.1.02.130 | Tehuacán-Cuicatlán | 4901.9 | 87 | 23 | 62 |
| 163 | 9.1.08.170 | Tiburón Ballena | 1459.9 | NA | NA | NA |
| 164 | 6.3.24.065 | Tula | 1 | 1 | 1 | 0 |
| 165 | 9.3.01.064 | Tulum | 6.6 | 0 | 0 | 9 |
| 166 | 3.7.01.018 | Tutuaca | 4369.9 | 62 | 14 | 37 |
| 167 | 9.7.02.116 | Uaymil | 891.2 | 0 | 0 | 14 |
| 168 | 1.7.02.058 | Valle de los Cirios | 25219.9 | 73 | 4 | 11 |
| 169 | 1.8.01.171 | Ventilas Hidrotermales de la Cuenca de Guaymas y de la Dorsal del Pac├¡fico Oriental | 1455.6 | NA | NA | NA |
| 170 | 5.3.02.006 | Volcán Nevado de Colima | 65.5 | 38 | 9 | 51 |
| 171 | 8.1.07.154 | Volcán Tacaná | 63.8 | 7 | 5 | 25 |
| 172 | 6.3.13.022 | Xicoténcatl | 8.5 | 4 | 2 | 48 |
| 173 | 8.4.03.135 | Yagul | 10.8 | 2 | 2 | 38 |
| 174 | 8.4.02.102 | Yaxchilán | 26.2 | 2 | 2 | 13 |
| 175 | 9.7.01.110 | Yum Balam | 1540.5 | 0 | 0 | 16 |
| 176 | 8.6.01.053 | Z.P.F. en los terrenos que se encuentran en los mpios. de La Concordia, Ángel Albino Corzo, Villa Flores y Jiquipilas | 1775.5 | 19 | 9 | 44 |
| 177 | 6.6.01.042 | Z.P.F.T.C.C. de los ríos Valle de Bravo, Malacatepec, Tilostoc y Temascaltepec | 1402.3 | 143 | 25 | 67 |
| 178 | 7.6.01.032 | Z.P.F.V. la Cuenca Hidrográfica del Río Necaxa | 421.3 | 20 | 8 | 56 |
| 179 | 5.1.04.163 | Zicuirán-Infiernillo | 2651.2 | 33 | 13 | 55 |
| 180 | 1.1.06.162 | Zona marina Bahía de los Ángeles, canales de Ballenas y de Salsipuedes | 3879.6 | 4 | 1 | ND |
| 181 | 1.3.06.161 | Zona marina del Archipiélago de EspÍritu Santo | 486.5 | NA | NA | NA |
| 182 | 1.3.05.156 | Zona marina del Archipiélago de San Lorenzo | 584.4 | NA | NA | NA |

**Supplementary Table 11. Area of three scenarios for conservation of Mesoamerican CWR in Mexico in federal protected areas and indigenous areas. For analysis, 20% of Mexico's terrestrial area is considered of each scenario.**

| **Scenario (20% area)** | **Area within PAs $km^2$( %)** | **Area within indigenous areas $km^2$ (%)** | **Area within PAs and indigenous areas $km^2$ (%)** |
|---|---|---|---|
| (a) all taxa | 42714.6 (10.5) | 165770.0 (40.8) | 19334.4 (4.8) |
| (b) taxa exclusively distributing in natural vegetation | 46060.8 (11.5) | 154905.9 (38.8) | 19861.9 (5.0) |
| (c) taxa associated to different habitats | 40170.3 (9.8) | 171405.4 (42.0) | 18237.9 (4.5) |

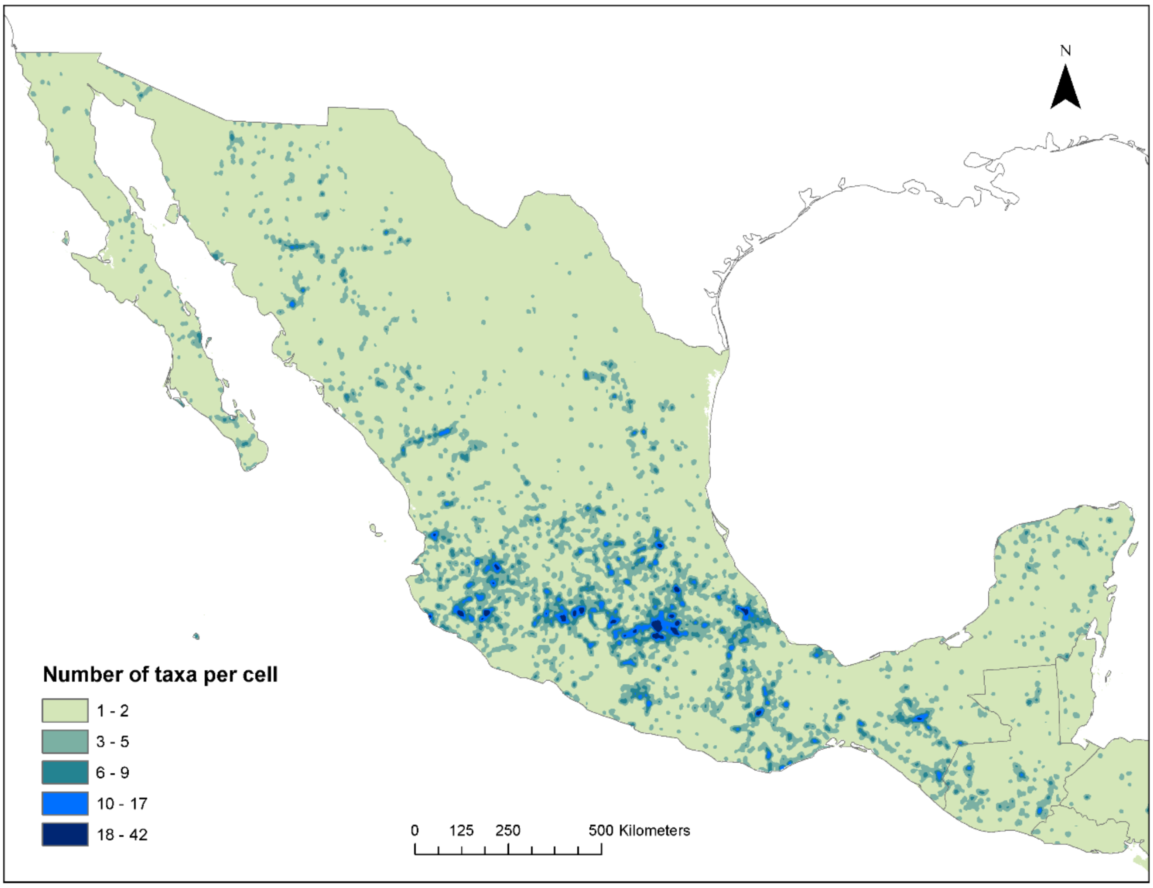

**Supplementary Fig. 1. Spatial pattern of taxa richness of selected Mesoamerican CWR in Mexico based on occurrence georeferenced data. Spatial resolution 5km$^2$.**

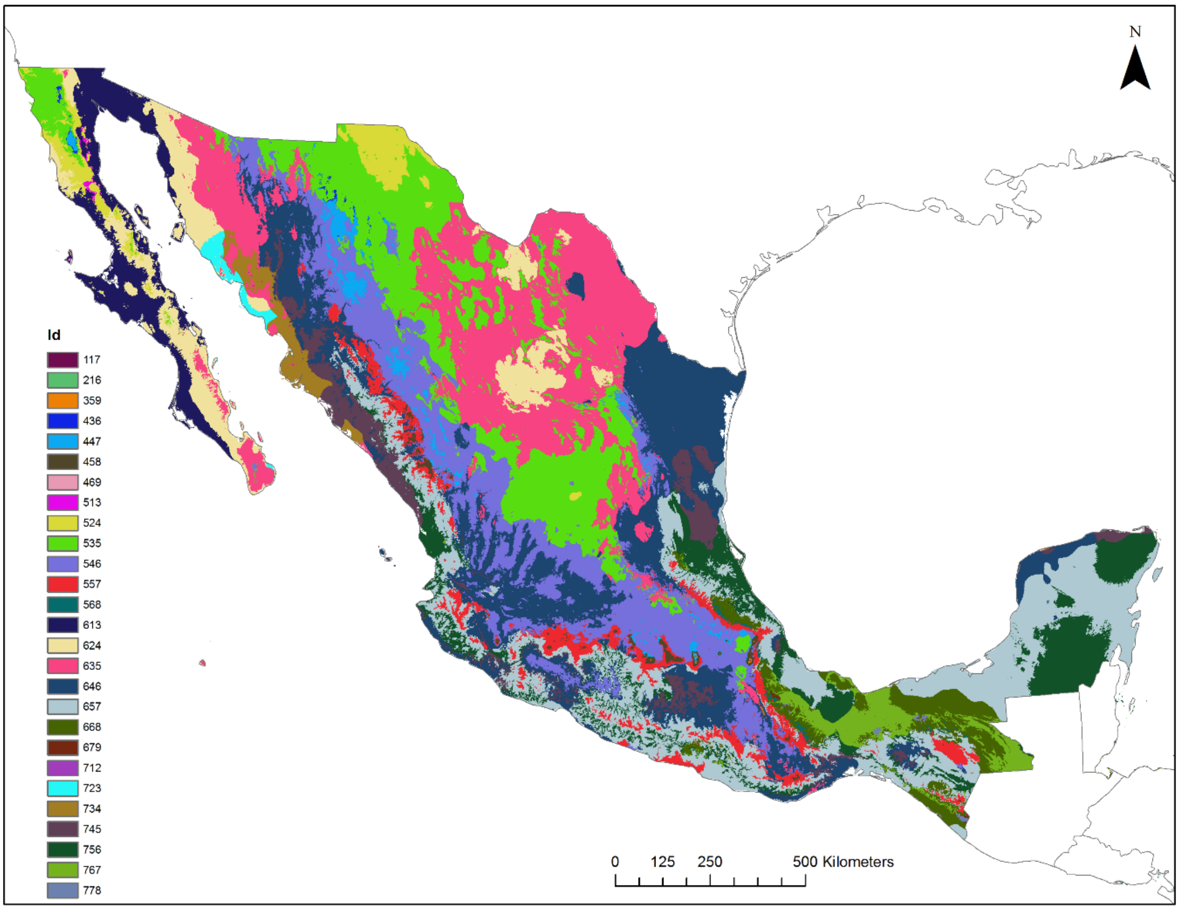

**Supplementary Fig. 2. Holdridge life zones of Mexico, based on biotemperature, annual precipitation and potential evapotranspiration ratio (see Supplementary Table 5). Life zones are represented by different colours (see code of numbers at Supplementary Table 5). Spatial resolution 1km$^2$.**

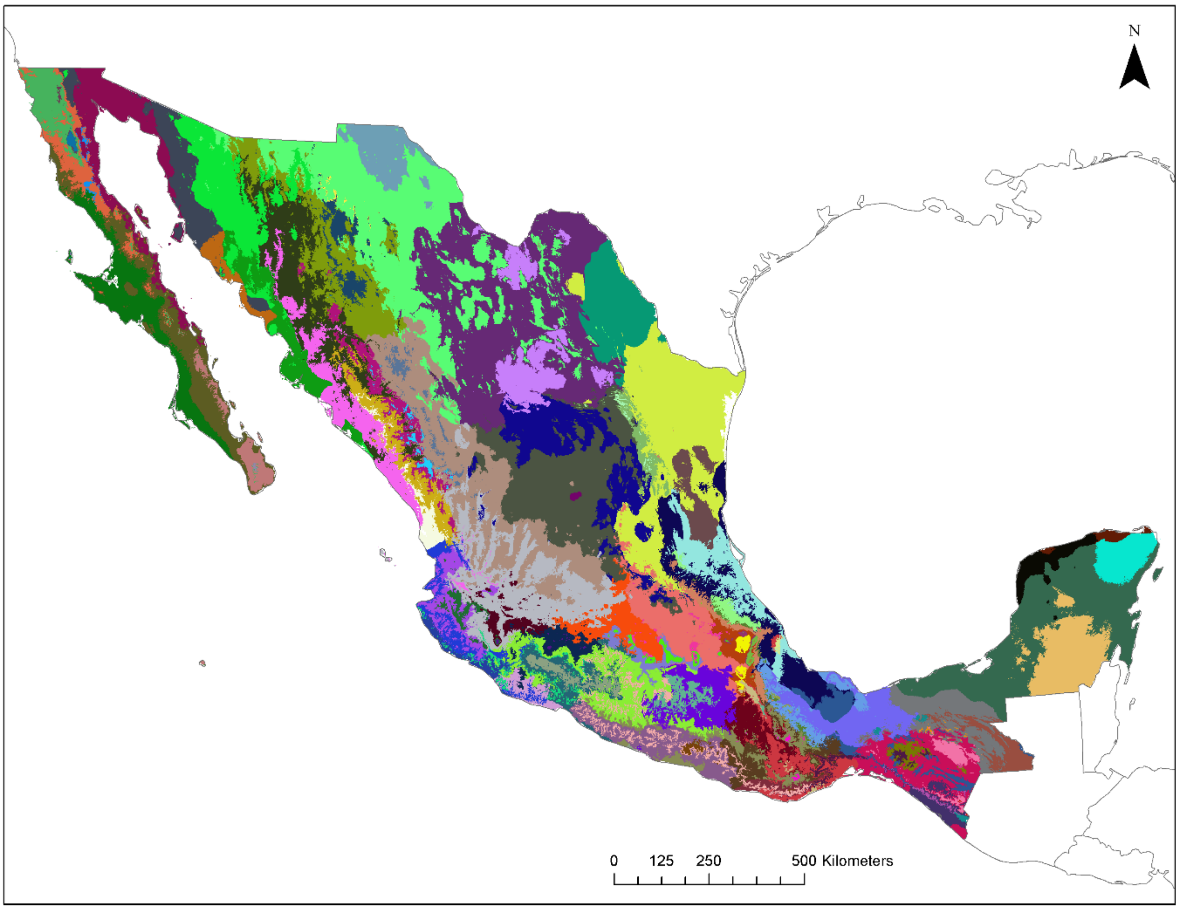

**Supplementary Fig. 3. Proxies of genetic diversity (PGD) for Mexico, based on environmental data (Supplementary Table 5, Supplementary Fig. 2) and historic drivers (Supplementary Table 6). The 102 PGD are represented by different colours. Spatial resolution 1km$^2$.**

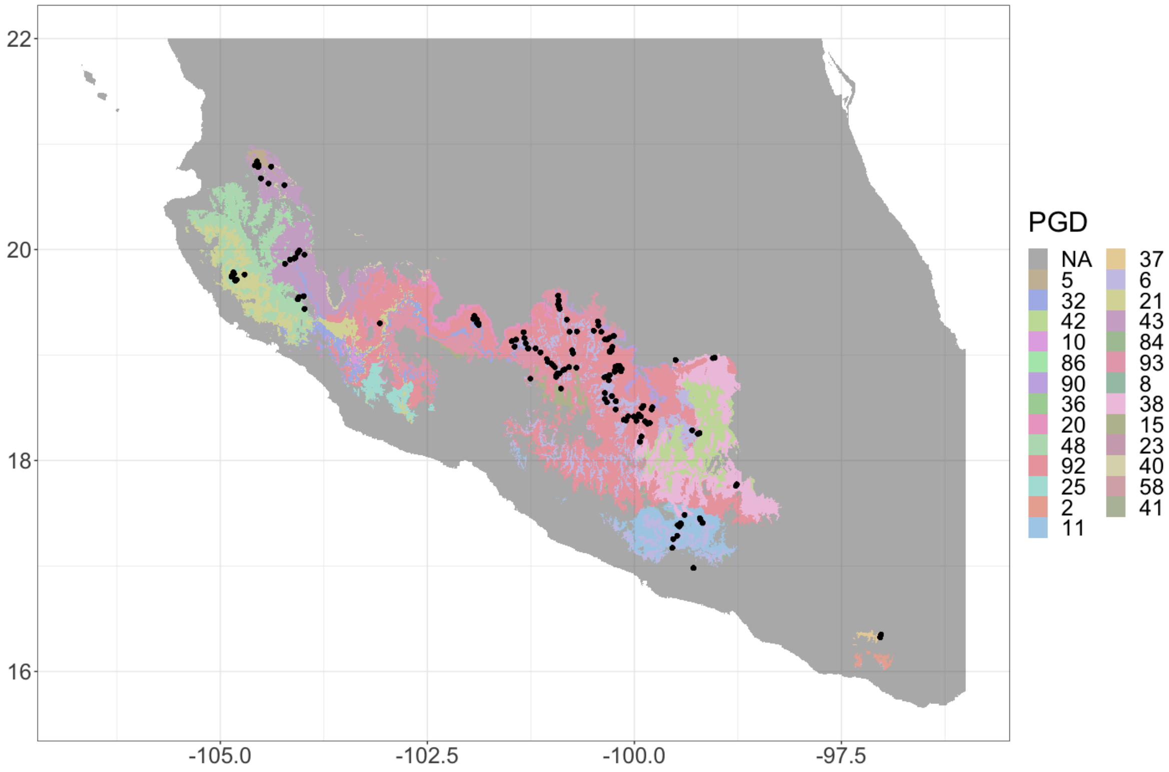

**Supplementary Fig. 4. Potential species distribution model of *Z. mays* ssp. *parviglumis* as given by PGD (background colours).**

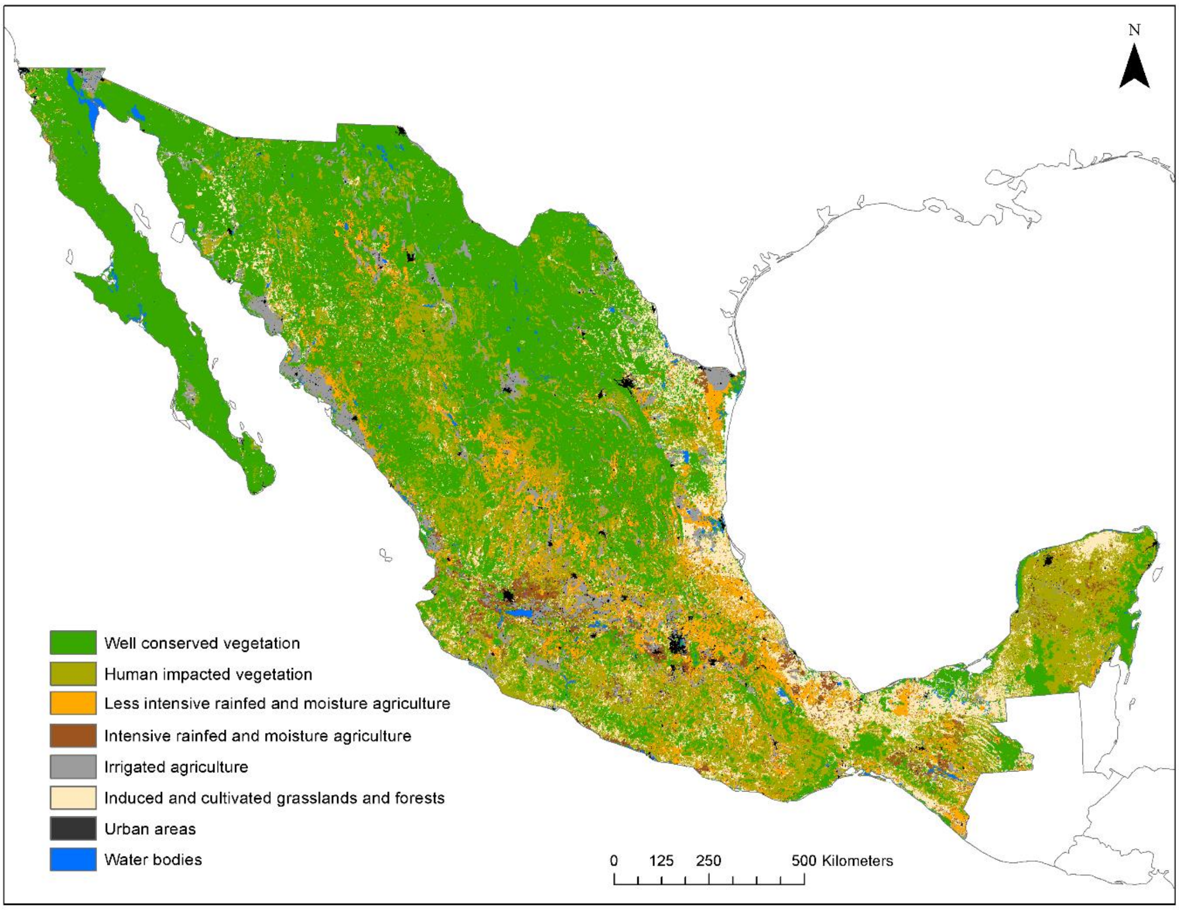

**Supplementary Fig. 5. Land cover map that was used to assess habitat preference of each taxon (Supplementary Table 7). Spatial resolution 1km$^2$.**

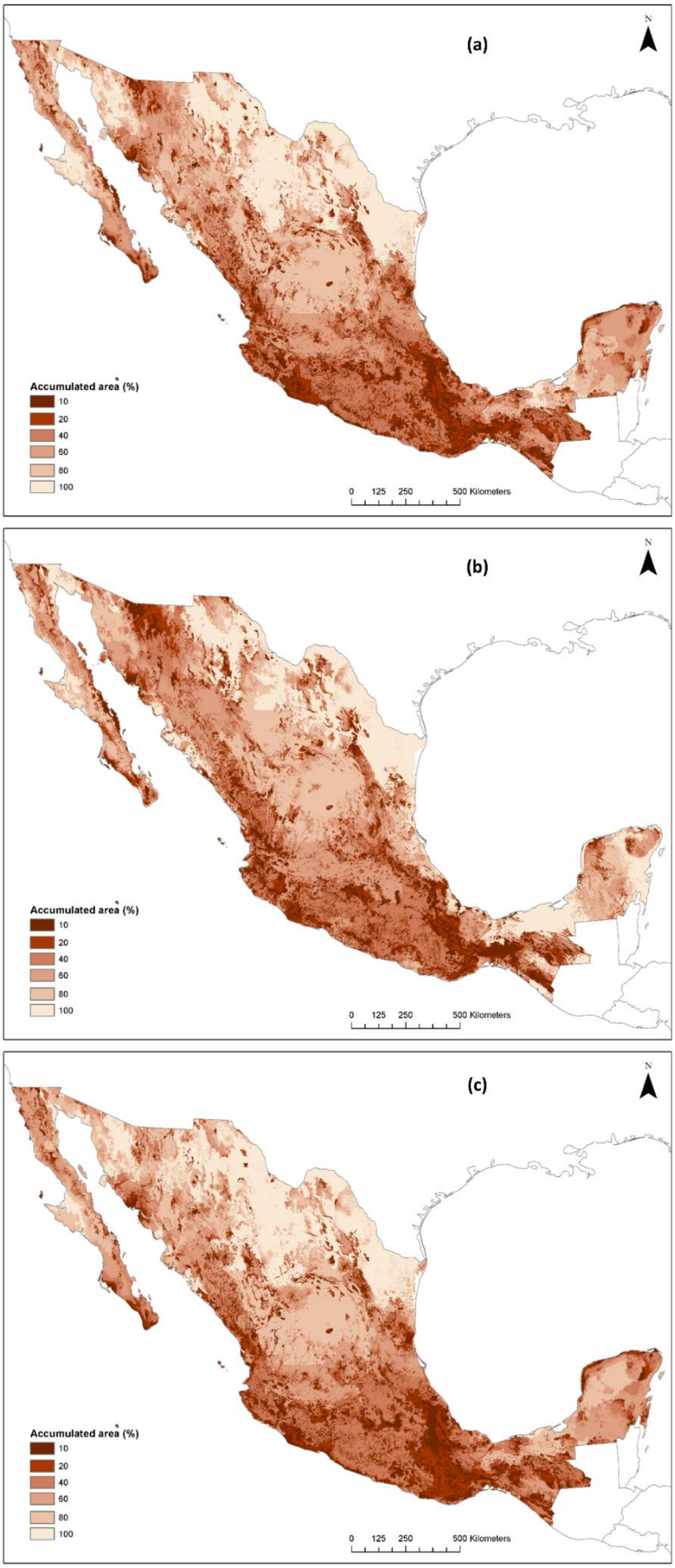

**Supplementary Fig. 6. Conservation areas for Mesoamerican crop wild relatives in Mexico, considering (a) all taxa, (b) taxa exclusively distributing in natural vegetation, and (c) taxa associated to different habitats. Spatial resolution 1km$^2$.**

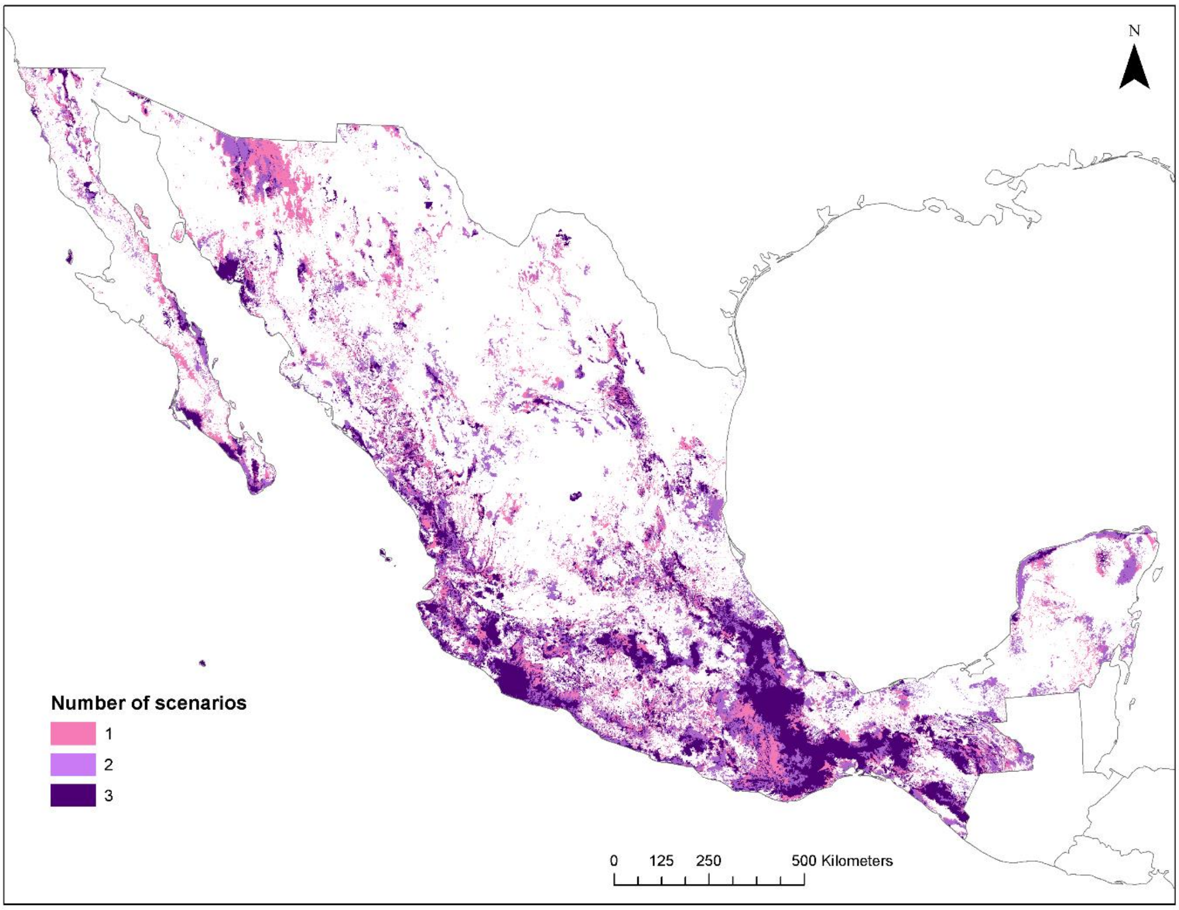

**Supplementary Fig. 7. Coincidence of three conservation scenarios, considering (a) all taxa; (b) taxa exclusively distributing in natural vegetation; and (c) taxa associated to different habitats. Twenty percent of Mexico's terrestrial area is highlighted of each scenario (see continuous values at Supplementary Fig. 6). Spatial resolution 1km$^2$.**

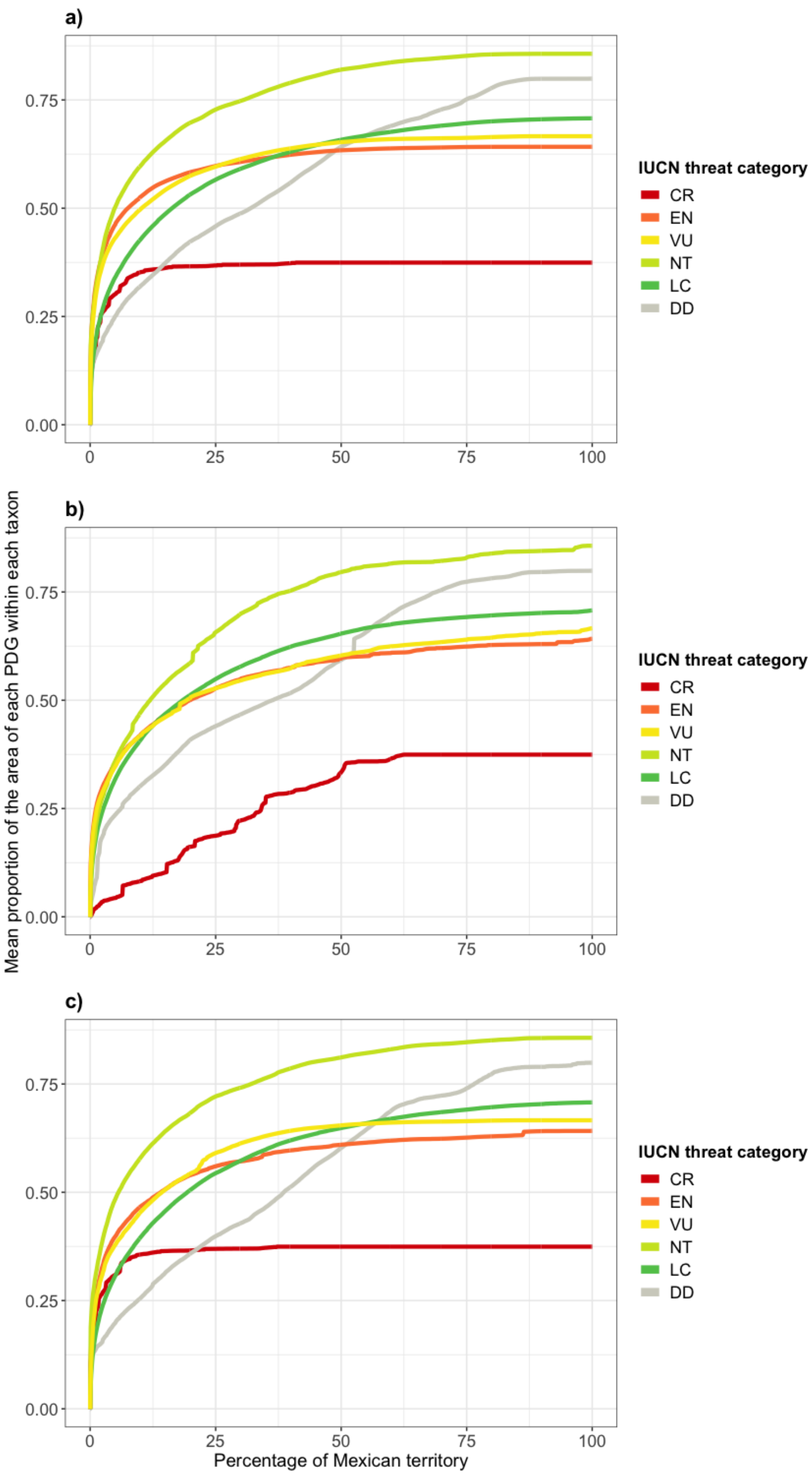

**Supplementary Fig. 8. Performance curves quantifying the proportion of taxa distribution ranges considered for each scenario, grouped by IUCN Red List Category. Scenarios considered (a) all taxa; (b) taxa exclusively distributing in natural vegetation; and (c) taxa associated to different habitats, i.e. natural vegetation, agricultural and urban areas.**

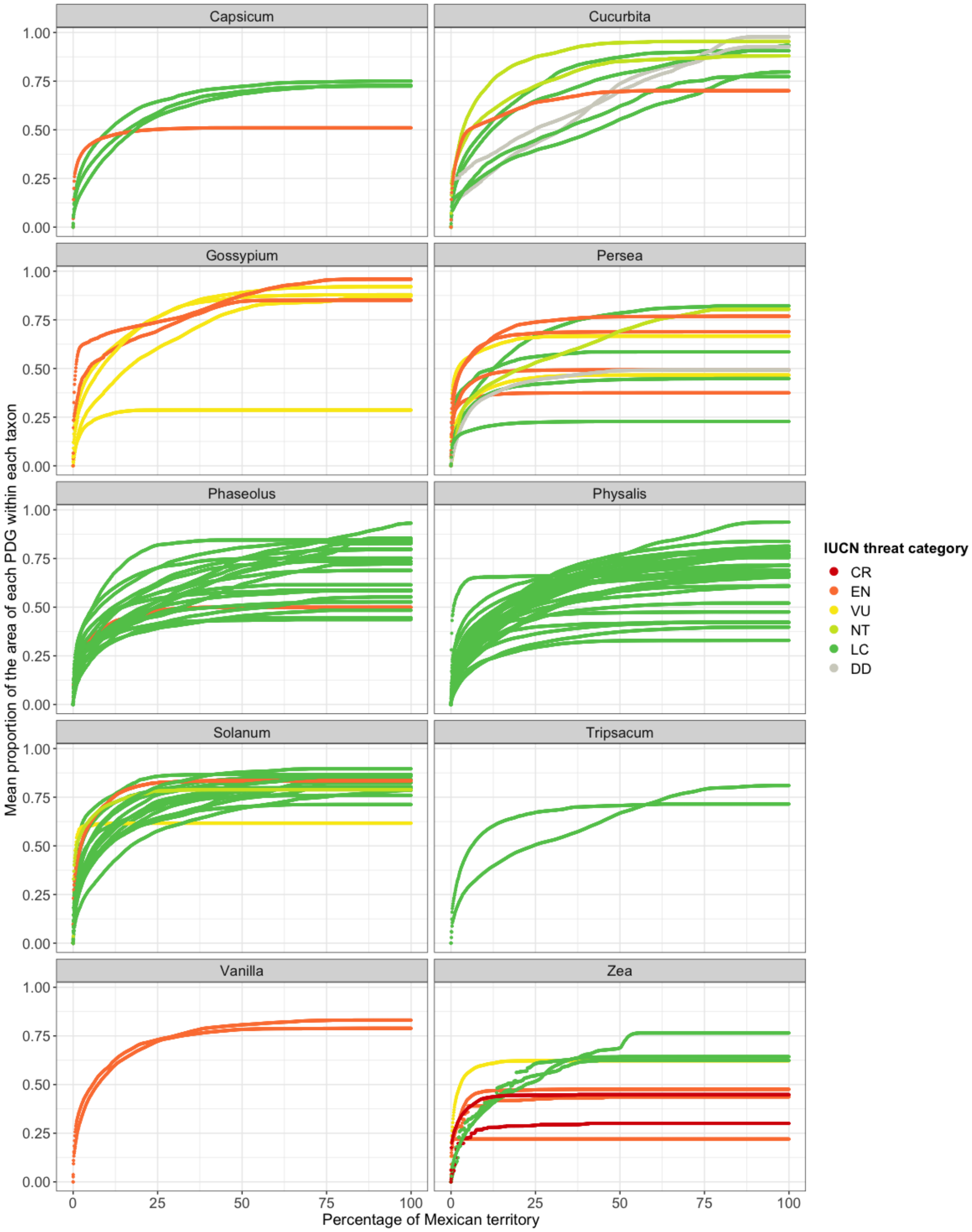

**Supplementary Fig. 9. Performance curves quantifying the proportion of taxa distribution ranges considering all priority taxa (see Fig. 5a or Supplementary Fig. 6a), grouped by genus; colours are according to its IUCN Red List Category.**

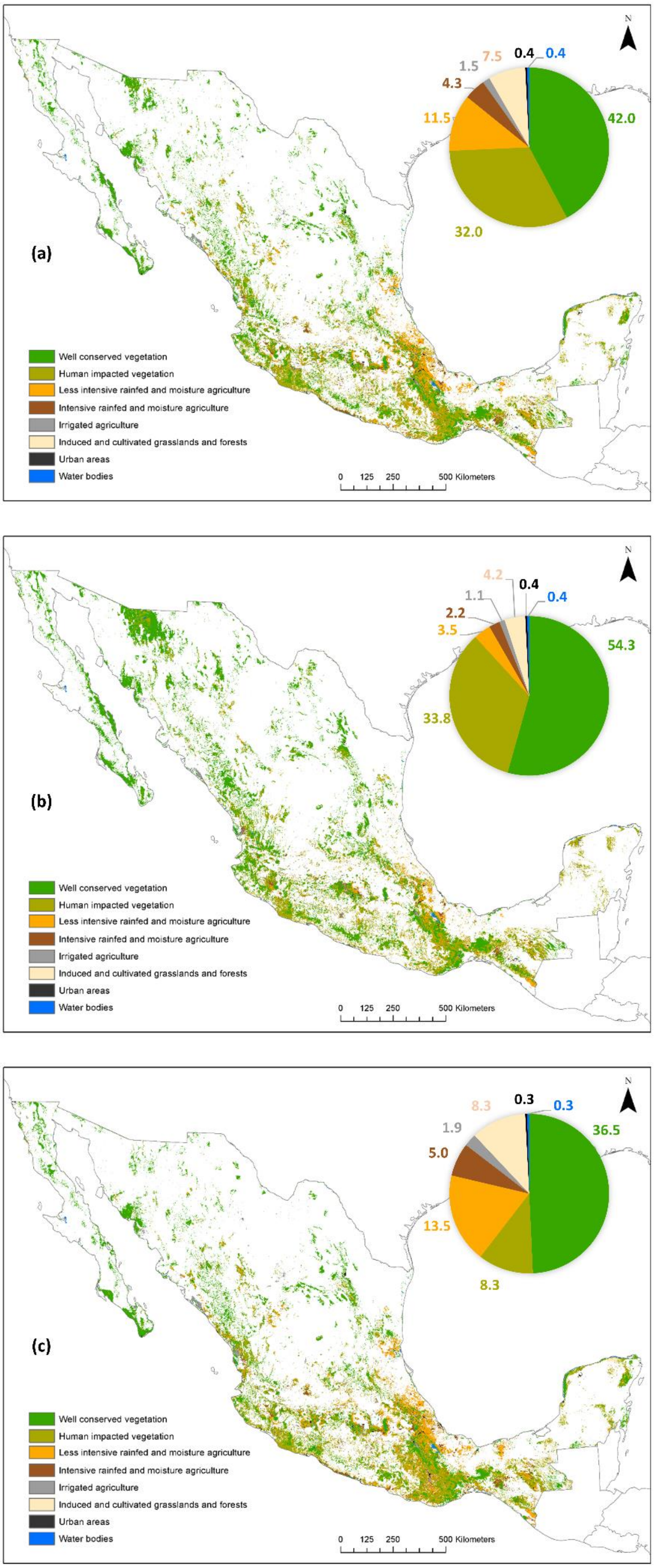

**Supplementary Fig. 10. Conservation areas of Mesoamerican crop wild relatives in Mexico according to land cover data used for the analysis (see Supplementary Fig. 5). Scenarios are based on proxies of genetic diversity, and consider (a) all taxa, (b) taxa exclusively distributing in natural vegetation, and (c) taxa associated to different habitats. Twenty percent of Mexico's terrestrial area is highlighted of each scenario.**

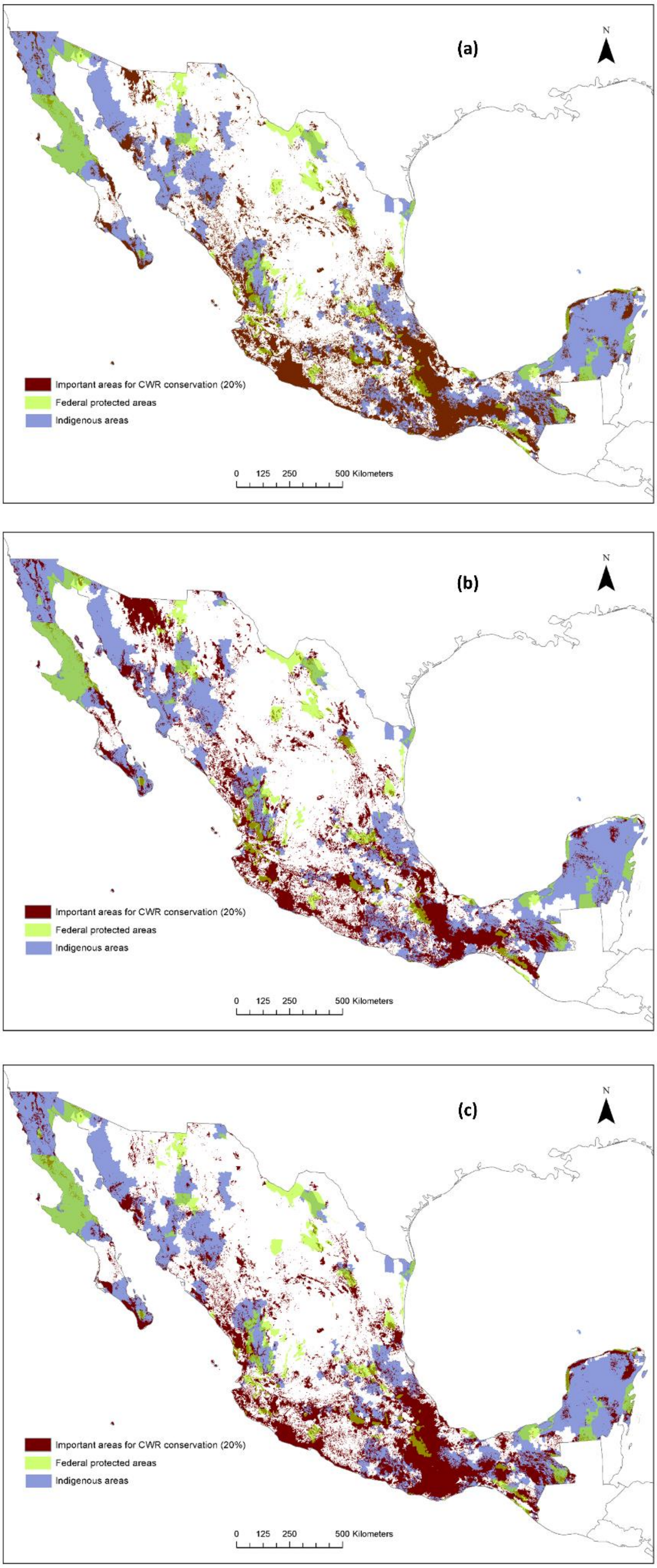

**Supplementary Fig. 11. Conservation areas for Mesoamerican crop wild relatives in Mexico, considering (a) all taxa, (b) taxa exclusively distributing in natural vegetation, and (c) taxa associated to different habitats, showing continuous values (left), and highlighting 20% of Mexico's terrestrial area, federal protected areas and indigenous areas. Spatial resolution 1km$^2$.**
